# Exploring the impact of lesser-known social dynamics on wolf populations through an individual-based approach

**DOI:** 10.1101/2020.01.24.918490

**Authors:** Sarah Bauduin, Oksana Grente, Nina Luisa Santostasi, Paolo Ciucci, Christophe Duchamp, Olivier Gimenez

**Affiliations:** Office Français de la Biodiversité, Juvignac, France; CEFE, Univ. Montpellier, CNRS, EPHE, IRD, Univ. Paul Valéry Montpellier 3, Montpellier, France; Office Français de la Biodiversité, Gap, France; Department of Biology and Biotechnologies, University of Rome La Sapienza, Viale dell’Università 32, 00185, Roma, Italy

**Keywords:** Gray wolf, Individual-based model, Pack dynamics, Population projection, R language

## Abstract

The occurrence of wolf populations in human-dominated landscapes is challenging worldwide because of conflicts with human activities. Modeling is an important tool to predict wolf dynamics and expansion, and help in decision making concerning management and conservation. However, some individual behaviors and pack dynamics of the wolf life cycle are still unclear to ecologists. Here we present an individual-based model (IBM) to project wolf populations while exploring the lesser-known processes of the wolf life cycle. IBMs are bottom-up models that simulate the fate of individuals interacting with each other, with population-level properties emerging from the individual-level simulations. IBMs are particularly adapted to represent social species such as the wolf that exhibits complex individual interactions. Our IBM predicts wolf demography including fine-scale individual behavior and pack dynamics based on up-to-date scientific literature. We explore four processes of the wolf life cycle whose consequences on population dynamics are still poorly understood: the pack dissolution following the loss of a breeder, the adoption of young dispersers by packs, the establishment of new packs through budding, and the different types of breeder replacement. While running different versions of the IBM to explore these processes, we also illustrate the modularity and flexibility of our model, an asset to model wolf populations experiencing different ecological and demographic conditions. The different parameterization of pack dissolution, territory establishment by budding, and breeder replacement processes influence the most the projections of wolf populations. As such, these processes require further field investigation to be better understood. The adoption process has a lesser impact on model predictions. Being coded in R to facilitate its understanding, we expect that our model will be used and further adapted by ecologists for their own specific applications.

## 1. Introduction

The gray wolf (*Canis lupus*) has been extirpated from most of the globe during the last century due to its competition with humans for wild prey, depredations on livestock and general persecution (Ripple et al., 2014). Most of the remaining populations were considered endangered in the early 20^th^ century (Mech and Boitani, 2003). However, numerous wolf populations are now under protection regimes and management actions favor species persistence or comeback (Chapron et al., 2014). Even though the presence of this large carnivore may play an important role in maintaining a healthy ecosystem and increase biodiversity, its recolonization is challenging. For example, the impact wolves exert on human activities such as livestock farming (Kaczensky, 1999; Lute et al., 2018), or the increasing threat of hybridization with dogs in human-dominated landscapes (Pilot et al., 2018; Randi, 2011; Randi et al., 2014) require an informed and effective management of the populations (Hindrikson et al., 2017). Management interventions may involve control of wolf populations through legal killings (Bradley et al., 2015; Harper et al., 2008; Santiago-Avila et al., 2018) or non-lethal management options (McManus et al., 2015; Treves et al., 2016) such as sterilization of breeders (Donfrancesco et al., 2019; Haight and Mech, 1997). In order to inform and help managers in making the best decisions, models are needed to forecast the impact of alternative management regimes on the population dynamics and viability of the species (Bull, 2006; Marescot et al., 2013). Not only can models help select the best management strategy among several, but they can also define the most effective application of any particular strategy (e.g., its intensity or frequency) (Haight and Mech, 1997). However, before predicting the impact of external factors on wolf populations, a good understanding of the species life cycle, as well as a reliable model simulating it, are required.

Different types of models have been used to project the dynamics of highly social species such as the gray wolf. Stage-structured models including age-, breeding- or dispersing-specific individual categories have been developed to predict population growth rate, and hence are relevant to make predictions at the population level (Haight and Mech, 1997; Marescot et al., 2012). Individual-based models (IBMs) have also been used to model population dynamics and have proven to be more flexible to represent species with complex social structure like wolves or coyotes (Chapron et al., 2016; Marucco and McIntire, 2010; Pitt et al., 2003). IBMs are bottom-up models that simulate the fate of individuals interacting with each other and/or their environment. IBMs can include many individual-level mechanisms (i.e., behavioral rules) and therefore can represent complex individual interactions as exhibited by these social species (Chapron et al., 2016; Haight et al., 2002; Marucco and McIntire, 2010; Pitt et al., 2003). Population-level results emerge from the individual-level simulations (Railsback and Grimm, 2012). IBMs are modular models, in that they are built as series of sub-models. Sub-models represent either processes of the species life cycle (e.g., reproduction, mortality) or external factors that modify the population structure (e.g., immigration, management). In this respect, IBMs can be very flexible as sub-models can be independently parameterized, reorganized or removed, or new ones can be added. This flexibility allows researchers to mimic the species life cycle very closely, to test different versions of the model by modifying only some sub-models, as well as testing the impact of external processes, such as different management actions, on simulated populations (Bull et al., 2009; Hradsky et al., 2019).

Researchers have used IBMs to simulate the impact of wolf-removal strategies on depredation and population viability (Haight et al., 2002), to test the robustness of abundance indices (Chapron et al., 2016) or to predict the recolonization of the species and the associated risk of depredation (Marucco and McIntire, 2010). The models were all based on the fundamental processes of mortality, reproduction and dispersal. They also enabled individuals to access to the breeder status by various means, such as pack creation or the replacement of a missing breeder in a pack. Additionally, Haight et al. (2002) and Chapron et al. (2016) included supplementary mortality processes mimicking different wolf removal strategies. However, other individual behaviors and social dynamics, not included in these IBMs, are known to occur in the wild. For example, Brainerd et al. (2008) found that the loss of breeders in a pack might disrupt its stability, depending on the pack size and number of remaining breeders, and may induce a pack dissolution. Specifically, small packs with only one breeder left had a high probability of breaking apart, and even higher when no breeder remained (Brainerd et al. 2008). In addition, several studies observed that when breeders died in packs, vacant male breeding positions were primarily filled by unrelated individuals, whereas vacant female breeding positions were mostly filled by subordinate females of the same packs, which were most likely the daughters of the former breeding females (Caniglia et al., 2014; Jedrzejewski et al., 2005; Vonholdt et al., 2008). These processes surely play a role in inbreeding avoidance. Another social process affecting wolf population dynamics is the adoption of unrelated individuals within packs. Young lone wolves not holding a territory sometimes join and are adopted, as subordinates, by packs (Mech and Boitani, 2003). Most adoptees are males of 1 to 3 years old (Meier et al., 1995; Messier, 1985). Adoptees seem to represent a non-negligible part of the populations, roughly estimated between 10% and 20% (Mech and Boitani, 2003). However, adoptees are generally not identified in wolf populations as it necessitates genetic sampling and relatedness analyses. The reasons behind this behaviour are still poorly known (Mech and Boitani, 2003). Finally, less common strategies of formation of new reproductive pairs through “budding” or “splitting” may influence the wolf establishment and reproduction dynamics. Budding is when a dispersing wolf pair with a mature subordinate from an existing pack and they establish a new pack of their own (Brainerd et al., 2008; Mech and Boitani, 2003). Pack splitting have been reported for large packs, mainly in North American wolf populations (Hayes and Harestad, 2000; Jedrzejewski et al., 2004; Meier et al., 1995; Vonholdt et al., 2008). A sub-group of individuals permanently splits off from their original pack to form a new one nearby, often due to the presence of two breeding pairs in the pack (Jedrzejewski et al., 2004; Mech and Boitani, 2003). It differs from budding in that no dispersing individual is involved in the process. Studies on wolf genetics, inbreeding (Caniglia et al., 2014; Vonholdt et al., 2008), hybridization (Fredrickson and Hedrick, 2006) or assessment of management alternatives (Haight et al., 2002; Haight and Mech, 1997) that fail to account for important processes of wolf social dynamics may provide limited or erroneous conclusions, leading to potential inappropriate management decisions. Unfortunately, the processes mentioned above are not often reported from field studies, they are rarely quantified and their details are poorly documented.

Here we develop an IBM to represent the wolf life cycle while exploring four lesser-known processes of its social dynamics, specifically: the pack dissolution following the loss of a breeder, the adoption by existing packs of young dispersers, the establishment of new packs through budding, and the different types of breeder replacement. Our model explicitly includes interactions between individual wolves, accounting for changes in wolf status (i.e., breeder vs subordinate, resident vs disperser), dispersal, and establishment processes while taking into account density-dependence and individuals’ relatedness. While we use up-to-date scientific literature to parametrize well-known individual behaviors and pack dynamics, we model multiple scenarios based on different parameters, similar to a sensitivity analysis, to explore the lesser-known processes. This also allow us to highlight the flexibility and modularity of our IBM. The variability among model predictions reveals processes that most affected wolf population dynamics, therefore indicating life-cycle traits that require further investigation to enhance reliability of population projections. We develop our model using the R language to facilitate its clarity, accessibility and uptake by ecologists. Given the flexibility of the model structure, our IBM can be easily parameterized according to different values, updated with improved knowledge on wolf dynamics, or modified to be adapted to other specific research or management questions on wolves.

## 2. Methods

### 2.1. Wolf IBM

#### 2.1.1. Main model structure

The model simulates the life cycle of the gray wolf using an individual-based structure, including fine-scale individual processes and pack dynamics through a non-spatially explicit approach. We calibrate the model for central European wolf populations (i.e., Alps, Apennines). These populations are recolonizing the territory, locally generating conflicts with human activities (Chapron et al., 2014). They have also been well monitored and we have good estimates for their demographic parameters (Marucco et al. 2009, Caniglia et al. 2014). The complete description of the IBM following the Overview, Design concepts, and Details protocol (“ODD” protocol) (Grimm et al., 2010, 2006) is provided in Appendix A.Simulated individuals represent wolves that are organized in packs. Each wolf holds a unique ID, a sex (male or female), an age, if it is a resident (i.e., member of a pack) or a disperser, if it attained breeder status or not, and a pack ID to which it belongs (if resident). The model also tracks each wolf’s genealogy and each individual has a mother ID and father ID, and a cohort ID (i.e., the year it was born). Wolves are individually aged as pups (1 year old), yearlings (2 years old), or adults (≥ 3 years old). We assume all wolves reach sexual maturity at 2 years old. We consider a pack when one or several individuals establish and become residents. A pack is usually composed by one breeding pair, potentially augmented by several non-breeding subordinates. Mortality causes (e.g., starvation, disease, vehicle collisions, culling, poaching, intraspecific strife) and rates differ among individuals, usually depending on their age (Marucco and McIntire, 2010) or their residence status (Blanco and Cortés, 2007). Moreover, higher population densities cause competition for food, space and mates, and may also induce a higher adult mortality due to intraspecific aggressions (Cubaynes et al., 2014). Following the death of one or both breeders, a pack can persist and breeders can be replaced. Wolves routinely disperse in response to competition and aggression related to food availability and breeding opportunity within their pack (Mech and Boitani, 2003). Non-breeding wolves are forced to leave the pack because of social drivers regulating group size within the territory (Ballard et al., 1987; Fritts and Mech, 1981; Fuller, 1989; Gese and Mech, 1991; Mech, 1987). In areas of high prey availability, dispersal is postponed (Ballard et al., 1987; Blanco and Cortés, 2007; Jimenez et al., 2017) and is rather triggered by the onset of sexual maturity of young wolves (Gese and Mech, 1991; Messier, 1985; Packard and Mech, 1980) so that most wolves have dispersed from their natal pack by the age of 3 (Gese and Mech, 1991; Jimenez et al., 2017). Given wolves dispersal abilities, individuals may move from one population to another through long distance dispersal (Blanco and Cortés, 2007; Ciucci et al., 2009) in immigration and emigration processes. In our model, “dispersers” or “dispersing individuals” include all non-resident individuals, comprising those that are actually dispersing (i.e., that left their natal pack and are dispersing through the landscape searching for an opportunity to establish a new territory and mate), as well as floaters (i.e., nomadic individuals without a territory and available either to replace missing breeders within packs, or to establish a new pack (Mancinelli et al., 2018)). One of the main processes for dispersing wolves to reproduce is to form a new pack with a dispersing mate of the opposite sex (Mech and Boitani, 2003). Dispersing individuals can also establish a new territory by themselves, waiting for a mate to later join them (Wabakken et al., 2001), therefore we also consider in our model that a solitary resident wolf holding a territory alone constitutes a pack. Although our IBM is not spatially explicit, we indirectly consider the relative spatial arrangement of wolves through their pack affiliation and the equilibrium density parameter (Table 1).

**Table 1:** Parameters used in the wolf IBM for the non-explored sub-models (see Figure 1). Probabilities and rates are estimates for a yearly time step.

| **Parameter** | **Sub-model in which the parameter is used (see Fig. 1)** | **Explanation** | **Value** | **Reference** |
| --- | --- | --- | --- | --- |
| Mean litter size | Reproduction | Number of pups that go out of the female den. Mean used for a Poisson distribution to generate the number of pups produced by a breeding pair. | 6.1 | Sidorovich et al. 2007 |
| Pup mortality | Mortality | Mortality probability for wolves in their first year of their life. | 0.602 | Smith et al. 2010 |
| Yearling mortality | Mortality | Mortality probability for non-dispersing yearlings. | 0.18 (sd = 0.04) | Marucco et al. 2009 |
| Non density-dependent | Mortality | Mortality probability for non-dispersing adults when | 0.18 (sd = 0.04) | Marucco et al. 2009 |
| adult mortality |  | the population is not at equilibrium density. |  |  |
| Density-dependent adult mortality | Mortality | Mortality probability for non-dispersing adults when the population is at equilibrium density.<br>Mortality probability is calculated using the density-dependent survival $\phi$ as:<br>$\text{logit}(\phi) = a + (b * ((\text{popDens} - c) / d))$<br>where popDens is the population density per 1000 km<sup>2</sup> | a = -1.196<br>b = -0.505<br>c = 53.833<br>d = 17.984 | Cubaynes et al. 2014 |
| Equilibrium density | Mortality;<br>Establishment in pairs;<br>Establishment by budding;<br>Establishment alone | Maximum number of packs the environment within which the population is simulated can hold. | 30 | Defined by the user |
| Territory size | Mortality | Average territory size (in km<sup>2</sup>) for wolves. Used to calculate the study area size with the equilibrium density<br>to estimate wolf density<br>through the model. | 104 | Mancinelli et al. 2018 |
| Mortality of<br>dispersing,<br>non-adopted<br>pups | Mortality | Mortality probability for<br>individuals that dispersed as<br>pups, following their pack<br>dissolution, and that were<br>not adopted by a pack<br>during the year. | 1 |  |
| Disperser<br>mortality | Mortality | Mortality probability for<br>dispersers (except<br>individuals that dispersed as<br>pups). | 0.31 | Blanco and<br>Cortes,<br>2007 |
| Dissolution<br>probability<br>for pack with<br>1 breeder left | Pack dissolution | Probability of dissolution<br>for small packs that have<br>only 1 breeding individual<br>left in the pack. | 0.258 | Brainerd et<br>al. 2008 |
| Dissolution<br>probability<br>for pack with<br>no breeders<br>left | Pack dissolution | Probability of dissolution<br>for small packs that do not<br>have any breeding<br>individual left in the pack. | 0.846 | Brainerd et<br>al. 2008 |
| Relatedness threshold | Replacement of breeding females by subordinates;<br>Establishment in pairs;<br>Establishment by budding;<br>Replacement of breeders by dispersers;<br>Replacement of breeding males by subordinates | Relatedness value between 1<sup>st</sup> cousins. Two individuals must have their relatedness coefficient less than or equal to this threshold to be considered unrelated. | 0.125 | Caniglia et al. 2014 |
| Mean pack size | Dispersal;<br>Adoption | Mean used for the Normal distribution to generate the sizes of each pack, representing their maximum pack size. | 4.405 (sd = 1.251) | Marucco and McIntire, 2010 |
| Pup dispersal probability | Dispersal | Relative probability (compared to the other age categories) for a pup to leave the pack and become disperser. | 0.25 | Haight and Mech, 1997 |
| Yearling dispersal probability | Dispersal | Relative probability (compared to the other age categories) for a yearling to leave the pack and become disperser. | 0.5 | Haight and Mech, 1997 |
| Adult dispersal probability | Dispersal | Relative probability (compared to the other age categories) for an adult to leave the pack and become disperser. | 0.9 | Haight and Mech, 1997 |
| Number of immigrants | Immigration/<br>Emigration | Number of immigrants arriving in the population.<br>All values have the same probabilities to be chosen. | 0, 1, or 2 | Defined by the user |
| Proportion of emigrants | Immigration/<br>Emigration | Proportion of dispersing individuals that emigrate outside of the study area. | 0.1 | Defined by the user |

The time step of the model is one year as the wolf life cycle is organized around reproduction that happens once a year. At each time step, all simulated individuals go through the same series of different sub-models representing different processes of the wolf life cycle, and each individual behaves differently according to its own characteristics. The first sub-models are, in order, reproduction, aging and mortality. They are followed by several sub-models related to changes in breeder/subordinate and resident/disperser status (Fig. 1). Most of the sub-models included in our IBM represent well-studied and well-quantified processes (Table 1). However, four processes of the wolf dynamics that we include are lesser known: the pack dissolution following the loss of a breeder, the adoption of young dispersing individuals by packs, the establishment of new packs using the budding strategy, and the different types of breeder replacement (Fig. 1). We explore different parameter values for the sub-models pack dissolution, adoption, and new pack establishment by budding (Table 2) and different timing (i.e., orders) for the sub-models concerning the breeder replacements (Fig. 1).

**Figure 1:**
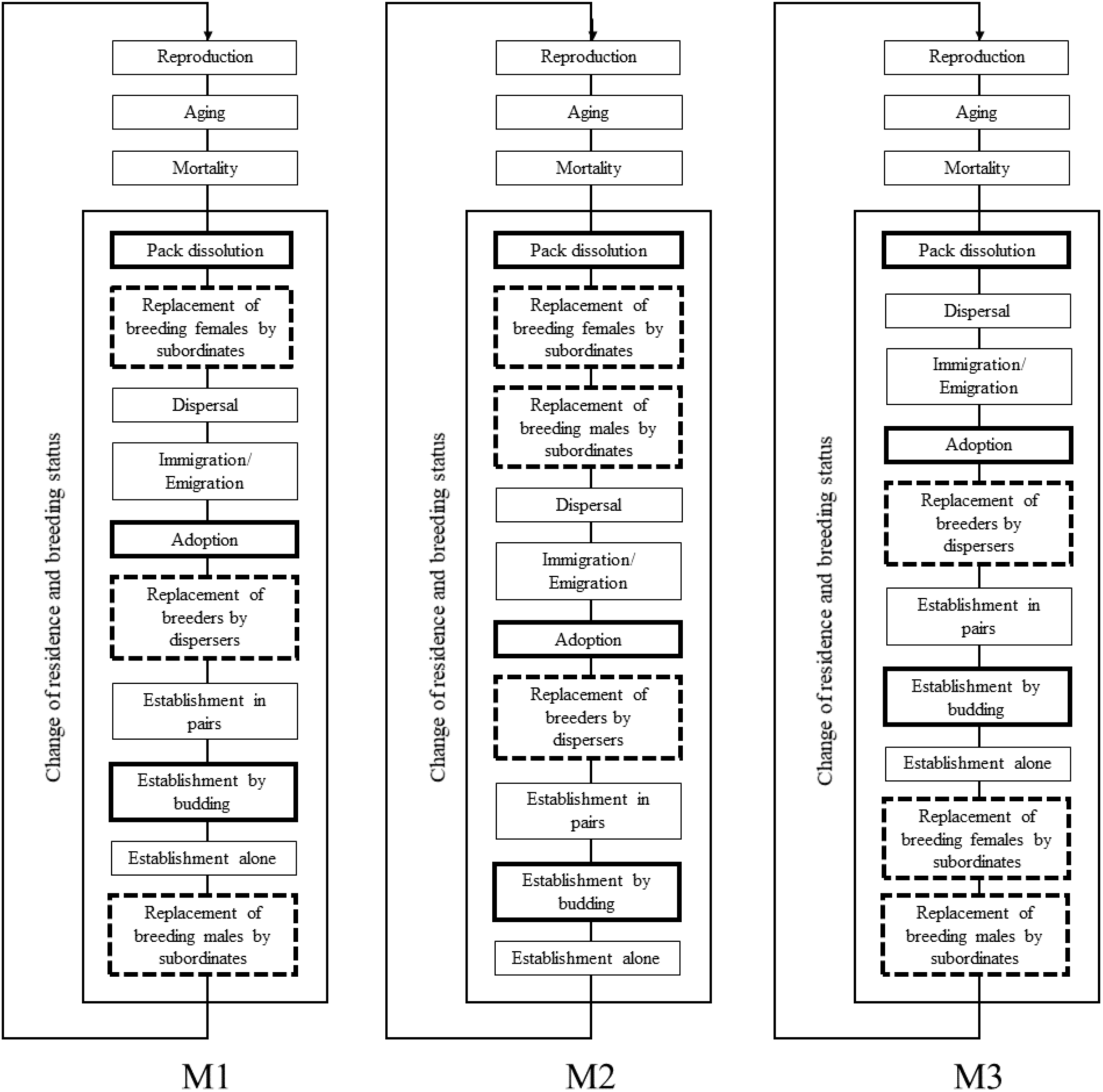
Diagram of three different versions of the wolf IBM (M1, M2 and M3). The sub-models in solid bold boxes are the lesser-known processes explored with different parameter values (Table 2). The sub-models in dashed bold boxes are the lesser-known processes related to breeder replacement for which their order, instead of their parameter, is tested with the three versions of the sub-models series: M1, M2, and M3. When a wolf population is simulated with one of these model versions, individuals go through each sub-model of the loop all together, one sub-model at a time, for as long as the simulation lasts. The loop of sub-models represents a one-year time step. (2-column fitting image)

**Table 2:** Parameters used in the wolf IBM for the explored sub-models (see Figure 1). Probabilities are estimates for a yearly time step.

| Parameter | Sub-model in which the parameter is used (see Fig. 1) | Explanation | Tested values |
| --- | --- | --- | --- |
| Pack size threshold for dissolution | Pack dissolution | Pack size to differentiate large and small packs. Packs with fewer individuals than this threshold are considered small and can dissolve if 1 or 0 breeding individual remains. | 3.1; 4.1; 5.1 |
| Probability of adopting | Adoption | Probability for a pack that has less member than its maximum pack size to adopt a young disperser. | 0.1; 0.5; 0.9 |
| Probability of budding | Establishment by budding | Probability of success for a disperser to establish a new pack by budding. This probability is multiplied by the density-dependent probability of establishment for the dispersers. | 0.1; 0.5; 0.9 |

#### 2.1.2. Initial population

A wolf initial population is needed to launch the IBM simulations. The user specifies the composition of its initial population and its attributes, namely the equilibrium density and immigration and emigration rates, to best represent the population he/she wants to model. Here is a simple example of the population we use in the following analyses. We build a fictive initial population of 10 packs and 5 dispersers, in an environment that can hold 30 packs in total (i.e., equilibrium density, Table 1). Specifically, the population comprises 5 packs of 2 breeders (5 years old each) with 2 pups (one male and one female); 3 packs of 2 breeders (5 years old each) with 1 yearling (male) and 1 pup (female); 2 packs of 2 breeders (5 years old each) with 1 adult (female, 3 years old); and 5 dispersers (3 females, 2 males, 2 years old each). We estimate the size of the area where the population is simulated as the number of packs at equilibrium density multiplied by the average territory size for wolf populations in the Apennines (104 km^2^) (Mancinelli et al., 2018). This area is used to calculate wolf density in density-dependent processes. We allow connections of the simulated population with other non-simulated wolf populations via an immigration of 0, 1 or 2 external wolves per year inside the simulated population, and an emigration of 10 % of the dispersing wolves from the simulated population outside of the study area. The parameters equilibrium density, number of immigrants arriving per year and proportion of dispersing wolves emigrating are randomly chosen. A modification of their values surely will change the projections of the population but the impact of these parameters on wolf populations are well understood and are therefore not explored further in this study.

#### 2.1.3. Well-known processes of the wolf life cycle

We define as well-known processes wolf dynamics that are well documented and well understood. These processes are usually included in wolf IBMs (Chapron et al., 2016; Haight et al., 2002; Marucco and McIntire, 2010). When coding the sub-models to reproduce these processes, we have reliable estimates to parameterize and time them in our model.

##### 2.1.3.1. Reproduction

We simulate that all packs with a breeding pair reproduce each year (Ciucci and Boitani 1999; Marucco and McIntire, 2010). We define the number of pups breeding pairs produce by sampling values in a Poisson distribution (Table 1) (Chapron et al., 2016). The sex of each pup is randomly chosen as male or female with a 1:1 ratio (Marucco and McIntire, 2010; Sidorovich et al., 2007). Newborn pups are of age 0. Pups are residents, with the pack ID of their parents, bear their mother and father IDs, and are assigned a cohort ID equal to the year of the simulation during which they are born.

##### 2.1.3.2. Aging

All individuals age one additional year in this sub-model. Pups of the year are now 1 year old, yearlings are 2 years old, and individuals of 3 years old enter the adult age class.

##### 2.1.3.3. Mortality

We simulate seven different mortality rates (Table 1), according to various combination of age and residence status of the individuals, and total number of packs related to the number at equilibrium density. Mortality is applied individually using a Bernoulli distribution. At each time step, the mortality probability is sampled from a Normal distribution using the mean and standard deviation parameters (Table 1) associated to the different categories of individuals.

###### 2.1.3.3.1. Mortality for non-dispersing individuals

We apply a different probability of mortality for non-dispersing individuals regarding their age category (i.e., pups, yearlings or adults) (Table 1). To mimic density-dependence in adult mortality (Cubaynes et al., 2014), we apply two mortality rates for this age category depending on the number of packs established in the population during any given year of the simulation. If the number of packs is below the equilibrium density, mortality is fixed (Table 1). If the number of packs reaches equilibrium density, we linked wolf mortality with wolf density following Cubaynes et al. (2014).

###### 2.1.3.3.2. Mortality for dispersing individuals

Dispersing yearlings in this sub-model are individuals whose pack dissolved in the previous year when they were pups and that were not adopted by any pack during that year. Their likelihood of survival alone is low without food or care from adults or yearlings (Brainerd et al., 2008; Mech and Boitani, 2003) so we set their mortality probability to 1. Otherwise, we set the same fixed mortality to all dispersing adults (Table 1).

###### 2.1.3.3.3. Mortality for old individuals

We do not model senescence or any increase in mortality rate with age. However, to represent a realistic age distribution in the population, we allow wolves to live up to their 15^th^ year of age (Marucco and McIntire, 2010), and all individuals of this age die entering the successive year of simulation.

##### 2.1.3.4. Dispersal

For each pack, and at each time step, we simulate the maximum number of individuals packs can support using a Normal distribution (Table 1). If a pack exceeds its simulated threshold, a certain number of individuals disperse until the number of wolves in the pack levels off at the threshold. While breeding individuals do not disperse, all other wolves can. Among these, the choice of dispersing individuals is based on relative probabilities related to their age category (Table 1).

##### 2.1.3.5. Immigration/Emigration

According to the immigration portion of the sub-model, at each time step, a determined number of immigrants enters the population (Table 1). Immigrants are simulated as dispersers, generated from another wolf population. Their sex is randomly chosen (i.e., male or female with a 1:1 ratio), and their age is simulated using a truncated Poisson distribution bounded between 1 and 15 with mean equal to 2 as dispersers are most commonly yearlings (Mech and Boitani, 2003). Immigrants do not belong to any pack of the simulated populations yet, and consequently are not breeders. As they were born outside the simulated population, they do not hold information about their mother ID and father ID (i.e., they are unrelated to any other individuals). However, once immigrating, they behave the same way (i.e., follow the same sub-model rules) as the native wolves. For the emigration portion of the sub-model, a proportion of the current dispersing individuals (Table 1), randomly chosen, leaves the simulated population. These individuals do not come back and their disappearance is similar to simulating their death.

##### 2.1.3.6. Pack establishment by breeding pairs

We define that reproductively mature male and female dispersers that are not closely related can pair bond, establish together as breeders, and form a new pack. The relatedness threshold chosen is that of the first cousins (Table 1); in wolves, breeding pairs are rarely more related than cousins, except when they have no other option (i.e., mating between siblings or parents and pups are generally avoided (Caniglia et al., 2014)). This relatedness threshold is the same for all sub-models. Establishment by breeding pairs is possible only if the number of existing packs has not yet reached equilibrium density. The density-dependent probability for dispersers to pair bond is defined by a Bernoulli distribution with probability equal to the number of packs that can be established until reaching equilibrium density divided by equilibrium density (i.e., the more packs there are, the less chance for dispersers to pair bond and establish new packs). Once a pair bond is established, both wolves become breeders and residents, sharing the same, new and unique pack ID.

##### 2.1.3.7. Pack establishment by single wolves

If the area is not at equilibrium density, our model allows mature dispersers that did not pair-bond to establish a territory by themselves. Similar to establishing in pair, the probability to establish a territory alone is density-dependent. Single wolves holding the new territory become breeders (even if no reproduction is possible yet) and residents of a new pack (i.e., they are assigned a new and unique pack ID). Then, at the next time step, a mature disperser of opposite sex will be able to take the vacant breeding position and finally reproduce.

#### 2.1.4. Lesser-known processes of the wolf life cycle

We define as lesser-known processes wolf dynamics that are known to occur in the wild but that are not extensively documented. These processes have not been included in previous IBMs for wolves (Chapron et al., 2016; Haight et al., 2002; Marucco and McIntire, 2010). When coding the sub-models to reproduce these processes, we do not have reliable estimates to parameterize or streamline them inside the life cycle available from the literature.

##### 2.1.4.1. Pack dissolution

We simulate that small packs whose social structure is impacted by the loss of one or both breeders may dissolve with different probabilities regarding how many breeders remain (Table 1). In the specific case where both breeders die and only pups remain in a pack, we consider that the pack always dissolves as pups are unlikely to maintain a territory by themselves. If these pups are not adopted during the current year, they die in the mortality sub-model during the next time step. When a pack dissolves, all former members of the pack become dispersers and do not belong to a pack anymore. Former breeding individuals also lose their breeder status. Brainerd et al. (2008) differentiate small packs, in which dissolution can occur following the loss of one or two breeders, from large packs in which dissolution never occur. They do not define a threshold between small and large packs but estimate that small packs have on average 2.36 individuals and large packs 5.75. We explore the importance of the pack dissolution process on the wolf population dynamics by varying this pack size threshold differentiating small packs from large ones (Table 2). A small threshold induces that only very small packs can dissolve, therefore minimizing the influence of pack dissolution on wolf dynamics. A large threshold allow more packs to be concerned, hence maximizing the influence of pack dissolution on wolf dynamics.

##### 2.1.4.2. Adoption

We define in our model that packs whose size is below their maximum threshold (estimated in the dispersal sub-model) can adopt as many individuals between 1 and 3 years old (inclusive) as allowed by their maximum pack size. Among potential adoptees, dispersing males are selected first. If there are no more males available to adopt, and packs are still small enough, then females are chosen next as adoptees. Once adopted, individuals become non-breeding (i.e., subordinate) residents and acquire their pack ID. Adoption has been observed (Mech and Boitani, 2003) but the rate at which this process occur is unknown. To explore how relevant this process is to affect wolf population dynamics, we defined different probability values for a pack to adopt (Table 2).

##### 2.1.4.3. Pack establishment by budding

Similar to the other strategies of establishment, budding is possible only if the number of packs in the population has not reached the equilibrium density. We define a density-dependent probability for a disperser to bud similar to the one of pack establishment through pair-bonding. Only mature dispersers can bud, and only with a non-breeding mature resident of the opposite sex that is not closely related (Table 1). Once budding occurs, both wolves that pair become breeders and residents, and obtain the same, new and unique pack ID. There are no detailed studies indicating how important budding is, compared to alternative ways of pack establishment. We explore the influence this process might have on wolf population dynamics by testing different probabilities of budding (Table 2) that we multiply to the density-dependence probability of pack establishment.

##### 2.1.4.5. Replacement of missing breeders

###### 2.1.4.5.1. Replacement of breeding females by subordinates

We select the subordinates to replace the missing breeding females by randomly choosing one of the subordinate mature females in the concerned packs. To mimic inbreeding avoidance in wolf packs, when then look at the relatedness between the chosen females and the current breeding males, if there is any. In case a breeding male is closely related to the female (Table 1), he may be replaced by a less related subordinate or a disperser (in sub-models *2.1.4.5.2. Replacement of breeding males by subordinates*, or *2.1.4.5.3. Replacement of breeders by dispersers*). We explore how the alternative sequence of the breeder replacement sub-models (i.e., breeders replaced by subordinates followed by breeders replaced by dispersers, or vice versa, independently for each sex; Fig. 1) affect projections of wolf population dynamics.

###### 2.1.4.5.2. Replacement of breeding males by subordinates

If there are several mature male subordinates in the pack where the male breeding position is vacant, the least related to the current breeding female, if there is any, is chosen to become breeder. If there are several unrelated subordinate males, or if there is no breeding female, one is selected randomly. In the case where the breeding female is related to the newly chosen breeding male but there is a mature female subordinate less related, the latter usurp the breeding female and the former breeding female is dismissed (i.e., becomes subordinate). If there are several unrelated mature female subordinates, one is selected randomly. We add that, in the particular case where a pack was not missing a breeding male but the breeding female obtained her breeding status during the time step (in *2.1.4.5.1. Replacement of breeding females by subordinates*) and she was too related to the current breeding male, one of the less related male subordinates can take over the male breeding position in this sub-model. All of these rules mimic inbreeding avoidance in wolves, except when there is no other choice to reproduce (Mech and Boitani, 2003). Once new breeding individuals are chosen, they will be able to mate the next year.

###### 2.1.4.5.3. Replacement of breeders by dispersers

Here, we simulate the replacement of all missing breeders, both females and males, by mature dispersers. First, missing breeding females are replaced by mature dispersing females, unrelated to the current breeding males in the packs if there is any. Then, missing breeding males are replaced similarly. Selected dispersers thus become breeders of packs, resident and acquire the ID of the pack they joined.

### 2.2. Model scenarios tested

We explore the impact of the four lesser-known processes on wolf population projections by varying and combining their different parameters values (Table 2) or timing (Fig. 1), resulting in 81 model scenarios (Appendix B). We run 200 replicates of each scenario for a 25-year simulation period, starting with the same initial population and same parameters for the well-known processes (Table 1).

#### 2.2.1. Model outputs

For each simulation, the complete population with the individual’s characteristics is available for each simulated year. The change in pack numbers is also recorded after each event potentially modifying their number: individual mortality (i.e., if all members of a pack die), pack dissolution, and the three types of new pack establishments. The model outputs we look at consist of six metrics expectedly crucial to evaluate wolf conservation and management. Specifically, we calculate, for each simulated year and for each replicate of each model scenario, the number of packs with a breeding pair. Then, we extract 1) at which year populations reach equilibrium density. This output represents the speed of the population expansion and is a key element in areas that are being recolonized by the wolf. We also calculate, after the last year of simulation, once all populations are at equilibrium density, for each replicate of each model scenario: 2) the number of packs with a breeding pair, as this corresponds to the reproductive potential of the population and is of importance for management issues related to population growth. 3) The number of packs newly established during the final year. This represents the pack turnover and the stability of the population that may affect mortality compensation, species expansion, or wolf-human conflicts (e.g., new packs may be more or less prone to attack livestock compared to old-established packs which know the associated risks and benefits). 4) The abundance of the population (i.e., total number of individuals) and 5) the proportion of residents and dispersers in the population. Population size is often required in management, and knowing the relative proportions of residents vs dispersers may help in understanding the demographic and social performance of the population and its potential to further expand its range. Finally, we compute 6) the relatedness between the two breeders in each pack. Inbreeding avoidance plays a big part in the wolf life cycle, affecting the replacement of the missing breeders and the creation of a new pack.

### 2.3. Sensitivity analysis

We run a sensitivity analysis on the parameters of the well-known processes (Table 1) to identify if some may influence the conclusions on the lesser-known processes. We use the model version M1 as a plausible model version, and the parameter values 4.1, 0.5, and 0.5 (i.e., all intermediate values) for the parameters of the sub-models pack dissolution, adoption and establishment by budding respectively. We run this model modifying the parameters of the well-known processes (Table 1) one parameter at a time by either increasing or decreasing its value by 5 % (Ovenden et al., 2019). We run 200 replicates of the model over 25 years to test each parameter. The model is considered sensitive to a parameter if a model output (i.e., mean value over the 200 replicates) with a modified parameter varies more than 20 % from the original results (Kramer-Schadt et al., 2005; Ovenden et al., 2019). We examine the six model outputs described above in the *2.2.1. Model outputs* section. We do not test the sensitivity of the model to standard deviation parameters (standard deviation of mortality and of pack size, Table 1). Regarding the density-dependent mortality function, we only test the sensitivity of the slope parameter and do not test the sensitivity to the intercept and to the parameters standardizing the population density (Table 1).

### 2.4. Model implementation

The IBM is coded in R 3.5.2 (R Core Team, 2014). We use the R package *NetLogoR* (Bauduin et al., 2019) to facilitate the implementation of the individual-based model structure in R language and the package *pedantics* (Morrissey, 2018) to calculate relatedness between individuals using their mother and father IDs. We use the packages *SciViews* (Grosjean, 2018) for the logarithms functions and *testthat* (Wickham et al., 2019) to implement tests in our model and verify the outcomes of the sub-models. The R files to run the model are available in the GitHub repository (https://github.com/SarahBauduin/appendix_wolfIBM) under the GNU General Public License v3.0.

## 3. Results

### 3.1. General results for the wolf IBM

Figures showing all model outputs for the 81 model scenarios are presented in Appendix C. All scenarios predict a growth of the wolf population. Starting from 10 packs and 43 wolves, after 25 years of simulation populations reach a mean of 29.4 (sd = 1.0) packs with a breeding pair, for an overall mean population size of 186.2 (sd = 12.4) wolves. In all model scenarios, populations stabilize and reach equilibrium density after about 10 years of simulation (mean = 9.8 years, sd = 2.8).

### 3.2. Effect of lesser-known social processes on wolf dynamics

We only present model outputs that show an impact from the parameterization or timing of the explored sub-models. Values reported are means and standard deviations calculated from all replicate simulations from each set of 27 model scenarios with the same parameter value or model version for the explored sub-model (i.e., 3 parameters or model versions * 27 model scenarios = 81 total).

#### 3.2.1. Pack dissolution

The different parameter values for the pack dissolution threshold influence the number of packs with a breeding pair and the total number of individuals in the population after the last year of simulation, as well as the number of new packs formed during the last year of simulation. Model scenarios where only small packs (i.e. with 3 individuals or less) may dissolve after breeders loss predict wolf populations with on average 29.6 packs with a breeding pair (sd = 0.8). Scenarios where larger packs may dissolve (i.e., packs up to 4 and up to 5 individuals) predict on average 29.4 packs (sd = 1.0), and 29.2 packs (sd = 1.1) respectively (Fig. 2.a). The most impacted model output by this sub-model parameterization is the number of new packs formed during the last year of simulation. More new packs are formed as larger packs are allowed to dissolve: with the threshold values 3.1, 4.1, and 5.1, the mean number of new packs created are equal to 1.3 (sd = 1.1), 2.2 (sd = 1.5) and 3.2 (sd = 1.7) respectively (Fig. 2.b). Regarding the number of individuals in the populations, the averages are equal to 189.4 (sd = 11.9), 186.3 (sd = 12.1), and 182.8 (sd = 12.3) respectively (Fig. 2.c).

**Figure 2:**
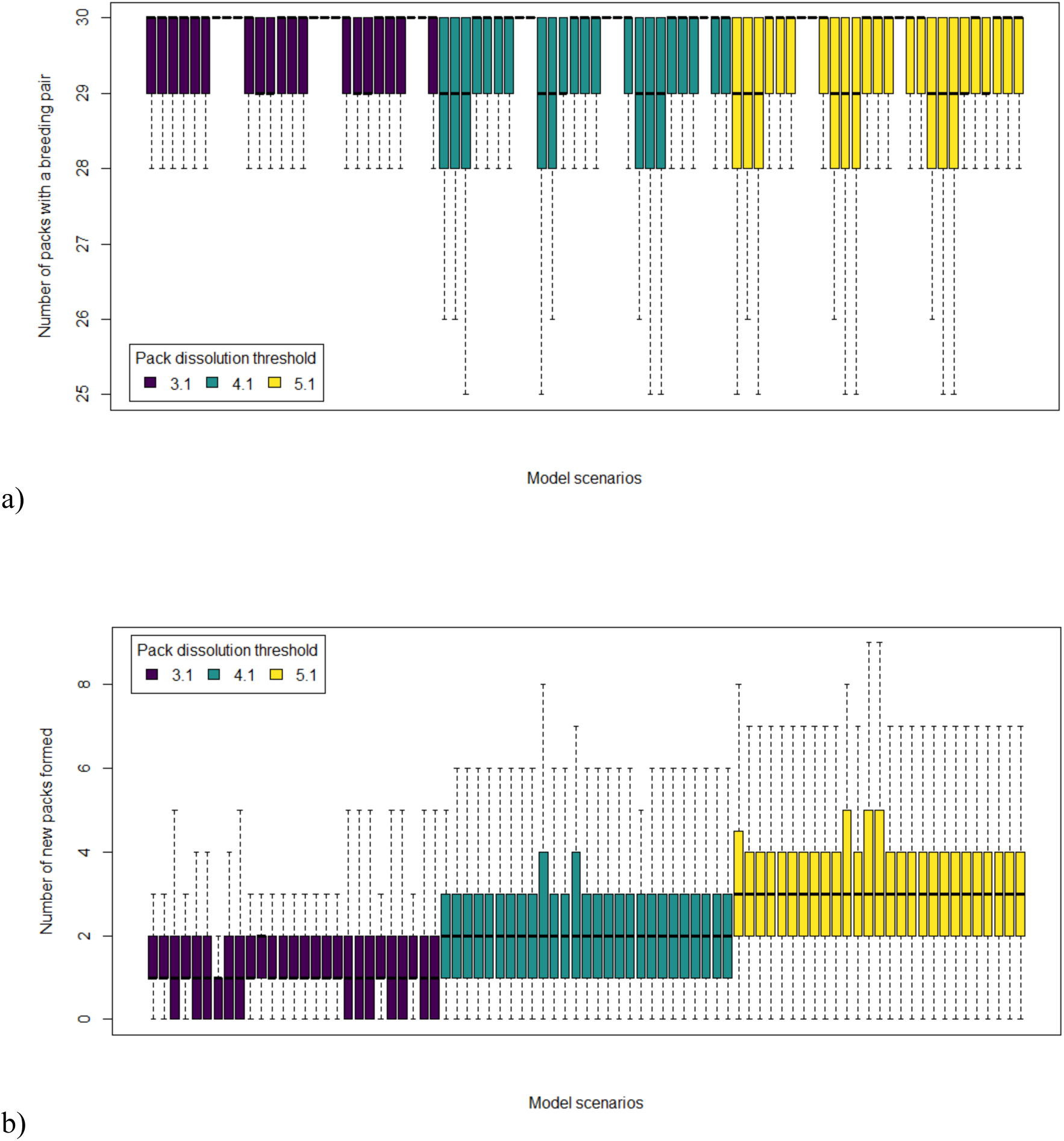
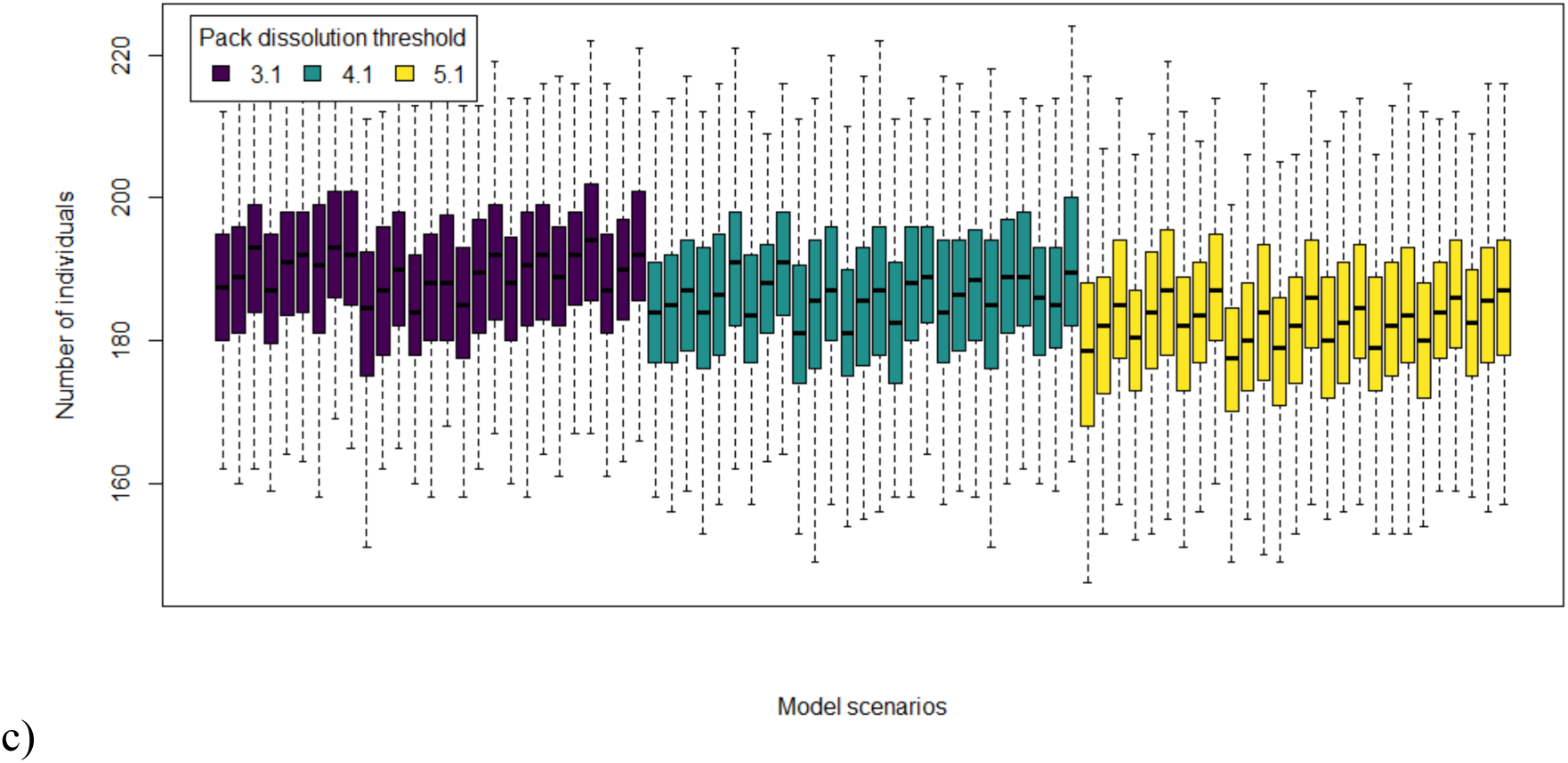
Model outputs influenced by the pack dissolution sub-model parameterization: a) number of packs with a breeding pair after the last year of simulation, b) number of new pack created during the last year of simulation, and c) total number of individuals in the population after the last year of simulation. Boxplots represent model output values extracted from the 200 replicates for each of the 81 model scenarios. They are color-coded and ranked according to the different values tested for the pack dissolution threshold (Table 2).

#### 3.2.2. Adoption

The different parameters for the adoption sub-model seem to only mildly influence the proportion of resident individuals in the populations. With an adoption probability set to 0.1, model scenarios predict on average 68.1 % of the population being resident (sd = 5.9 %), with an adoption probability equal to 0.5, the result is of 69.7 % (sd = 5.4 %) and with an adoption probability equal to 0.9, the result if of 71.2 % (sd = 5.8 %) (Fig. 3). The impact of varying the adoption probability from 0.1 (i.e., almost no adoption occurring) to 0.9 (i.e., adoption happening very often when possible) is very low on this model output and non-existent on the other ones (Appendix C).

**Figure 3:**
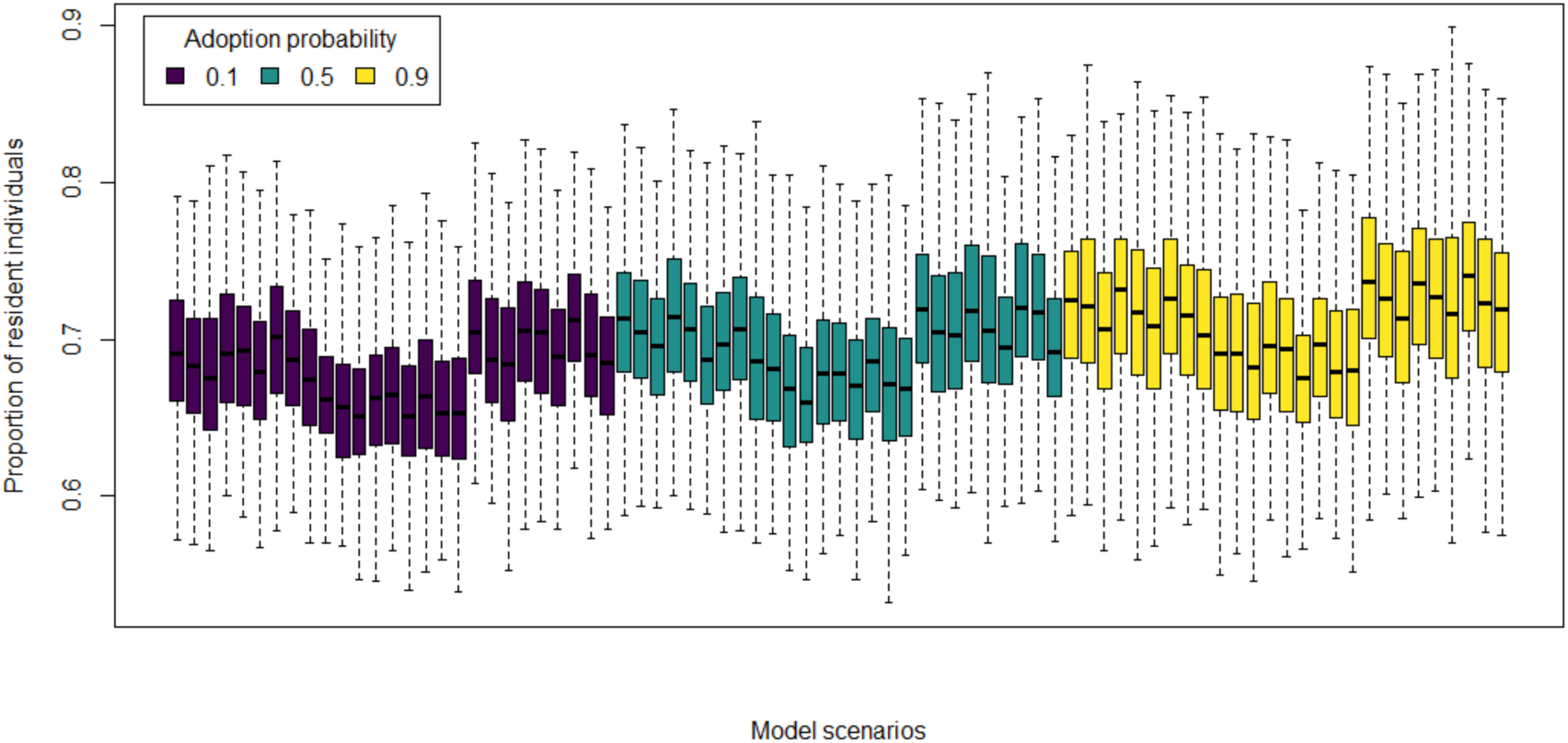
Model output influenced by the adoption sub-model parameterization: proportion of resident individuals in the population after the last year of simulation. Boxplots represent model output values extracted from the 200 replicates for each of the 81 model scenarios. They are color-coded and ranked according to the different values tested for the adoption probability (Table 2).

#### 3.2.3. Pack establishment by budding

The model outputs the most impacted by the different parameter values for the probability of establishment by budding are the time at which populations reach equilibrium density and the number of packs with a breeding pair at the end of the simulation. The lowest probability of budding (i.e., equal to 0.1) project wolf populations reaching the equilibrium density the latest, on average after 11.8 years of simulation (sd = 3.0) (Fig. 4). With the budding probability equal to 0.5, populations reached that point on average after 9.6 years (sd = 2.2) (Fig. 4). With the highest probability of budding (i.e., equal to 0.9), populations reach the equilibrium density the fastest, on average after 8.2 years of simulation (sd = 1.7) (Fig. 4). The observed differences in numbers of packs after the last year of simulation follow the same patterns: the highest budding probability produced the highest mean values. Model scenarios with a budding probability equal to 0.1 predict wolf populations with on average 29.0 packs (sd = 1.2), with a probability of 0.5, there are on average 29.4 packs (sd = 1.0), and with the budding probability equal to 0.9, there are on average 29.7 packs (sd = 0.6).

**Figure 4:**
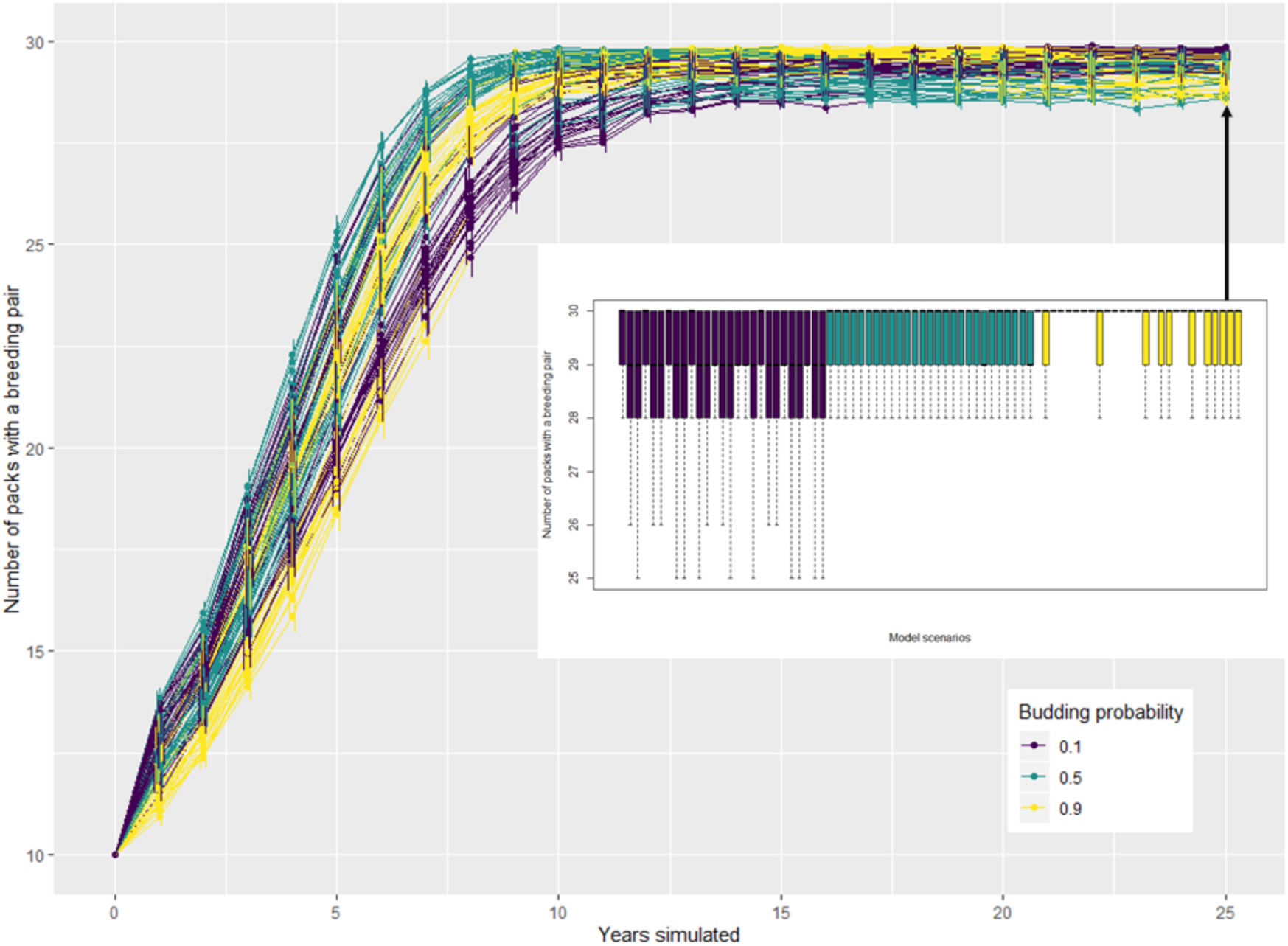
Model outputs influenced by the pack establishment by budding sub-model parameterization: time at which populations reach equilibrium density and number of packs with a breeding pair after the last year of simulation (figure inset). Line figure represent the mean values and their 95 % confidence intervals, per year, from the 200 replicates for each of the 81 model scenarios. Boxplots represent model output values extracted after the last year of simulation from the 200 replicates for each of the 81 model scenarios. Lines and boxplots are color-coded (and ranked for boxplots only) according to the different values tested for the probability of establishment by budding (Table 2).

#### 3.2.4. Replacement of missing breeders

The model versions testing the different timing of sub-models concerning breeder replacement influence mildly the proportion of resident individuals in the population after the last year of simulation, but it greatly impact the relatedness value between male and female in breeding pairs. Model versions M1 and M3 have similar results, and they both differ compared to those obtained using M2. M1 (Fig. 1) with the replacement of missing breeding females by subordinates first and the replacement of missing breeding males by dispersers first, predict populations with a mean proportion of resident equal to 70.3 % (sd = 5.3 %) (Fig. 5.a). It is similar to M3 (Fig. 1) where the replacement of missing breeders was done primarily by dispersers for both sexes; predicted populations have on average 71.2 % of resident individuals (sd = 5.5. %) (Fig. 5.a). In M2 (Fig. 1) where the replacement of missing breeders was done primarily by subordinates, predicted populations have on average 67.5 % of resident individuals (sd = 5.1 %) (Fig. 5.a). The influence of the different model versions is greater on the relatedness between breeders. For M1, the mean relatedness is equal to 0.06 (sd = 0.03), similarly as for M3 (mean = 0.06, sd = 0.04). For M2, it is equal to 0.26 (sd = 0.31) (Fig. 5.b). M1 and M3 favor the replacement of at least one missing breeder by a disperser and keep relatedness between breeders very low. In M2, missing breeders are primarily replaced by subordinates and mating between related individuals frequently occur.

**Figure 5:**
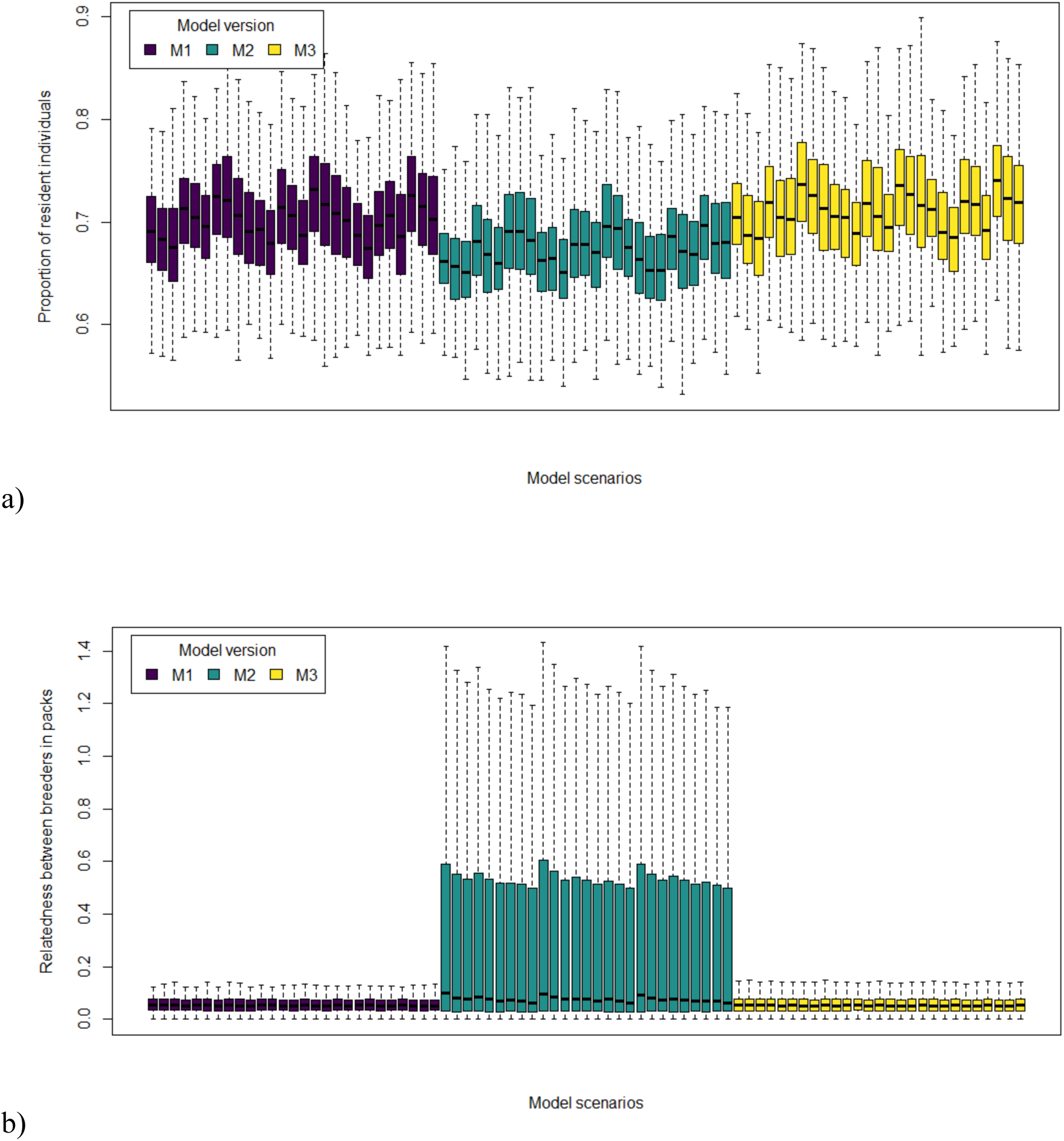
Model outputs influenced by the timing of the different sub-models simulating the replacement of missing breeders: a) proportion of resident individuals in the population after the last year of simulation, and b) relatedness value between the male and female in breeding pairs after the last year of simulation. Boxplots represent model output values extracted from the 200 replicates for each of the 81 model scenarios. They are color-coded and ranked according to the different model versions tested (Fig. 1).

### 3.3. Sensitivity analysis

As expected, the parameter affecting the most model outputs, apart from the ones tested for the lesser-known processes, is the equilibrium density. The number of new packs created is sensitive to this parameter. All the other model outputs are slightly sensitive to the equilibrium density parameter but not in the range of [- 20 %; + 20 %]. Pup mortality is the second parameter affecting the most model outputs but none vary more than the range of [- 20 %; + 20 %]. Overall, the six model outputs we look at are not sensitive to the parameters of the well-known processes (Table 1). The complete table with the value tested for the parameters and the results for the model outputs is available in Appendix D.

## 4. Discussion

We developed an IBM to represent wolf demography and pack dynamics while exploring lesser-known processes of the species social dynamics. We explored different parameterization and timing for the processes of pack dissolution, adoption, establishment by budding and replacement of missing breeders. The predictions from the different model scenarios pointed out the importance of the pack dissolution, establishment by budding and replacement of missing breeders processes in wolf population projections. Further research is needed to better understand those and the mechanisms behind, to be able to correctly incorporate them in the IBM with reliable parameter values and timing among the different sub-models. The different relative importance given to the adoption process did not modify much the model outputs, possibly indicating a lesser influence of this process on wolf dynamics. Our model also innovatively accounted for relatedness between individuals and density-dependence. Adding a genetic component to our model is especially important to investigate hybridization and inbreeding depression that can be of great concern for wolf management and conservation (Bohling and Wairs, 2015). Our analyses also highlighted the modularity and flexibility of our IBM that can be adapted and used by ecologists to explore various questions and test different hypotheses on wolf ecology, management, and conservation.

To explore the process of pack dissolution, we tested different values for the pack dissolution threshold to allow more or less packs to dissolve, given different importance to this process in the wolf dynamics. Diminishing the importance of pack dissolution produced populations often being at equilibrium density with few new packs created, mimicking a very stable population. The other way a pack can disappear, except than by dissolution, is by the death of all its members at once, which rarely happened in the simulations. On the contrary, allowing more packs to dissolve reduced the number of packs, freed space and partners to create new packs, and therefore favored the pack turnover. Still, the creation of new packs did not seem to reach the same intensity as the one of pack loss, as the final number of packs and individuals were slightly lower when pack dissolution was important. However, pack dissolution influenced more the composition of the packs at the individual level (i.e., pack turnover impacted) than the predictions at the population level (e.g., total number of packs and individuals). This is in agreement with a study by Borg et al. (2015) showing that with or without pack breaking down following breeder losses, the overall dynamics of the population was quite similar (e.g., number of packs), highlighting a compensation mechanisms. On the other hand, population stability is a key element in social species where individual personalities and group compositions matter, such as regarding hybridization (Bohling and Wairs 2015) and depredation (Allen, 2014), as well as their associated management actions. This process requires more field investigations to better understand the conditions leading to pack dissolution.

Budding is a strategy to establish and create new packs. With a low probability to bud, less packs are created and, as expected, simulations with the smallest probability of budding projected populations which were the slowest to reach equilibrium density. Inversely, maximizing the importance of budding projected populations reaching equilibrium density the fastest. At the end of the simulation, when the populations were at equilibrium density, the same pattern occurred with the more the budding strategy was used, the more packs were present in the populations. However, the different budding probability only slightly influenced the number of packs when the population was at equilibrium density. A better understanding of this process seems relevant to understand its relative importance regarding the other strategies of establishment to produce reliable predictions, even more during the growing phase of the population.

Having the replacement of the missing breeding female by a subordinate before a disperser (M1) has been documented in some study sites (Caniglia et al., 2014; Jedrzejewski et al., 2005; Vonholdt et al., 2008) but the regularity of this behavior is debated. The modified model version where the replacement of the missing female breeder was done by a disperser before (M3) produced very similar model outputs. However, the model version where both missing breeders were replaced by subordinates first (M2) produced very different predictions for the relatedness between breeders, and to a lesser extent on the proportion of resident individuals in the population. If we consider a breeding pair related when their relatedness coefficient is larger than 0.125 (Caniglia et al., 2014), only 0.8 % of the breeding pairs were related in populations predicted with model versions M1 and M3. In contrast, 38.1 % of the breeding pairs were related in populations predicted by M2. Vonholdt et al. (2008) evaluated that 7 % of the breeding pairs in the Yellowstone grey wolf population were related. The difference in our predictions and the value found by Vonholdt et al. can be due to spatiality. In our model versions M1 and M3, all dispersers are available to replace missing breeders, hence reducing the risk of inbreeding. In the wild, dispersers can be too far away and therefore not available to the packs missing breeders, hence inducing a replacement of the missing breeders by subordinates and inbreeding. Understanding the functioning of the different types of breeder replacement and their timing is crucial seeing their impact on model predictions. Studies on wolf genetics could conclude completely differently if using one or another model version, potentially inducing detrimental management recommendations for small isolated populations if predictions were not reliable. The modular and flexible construction of our IBM model can allow future users to organize the replacement of the missing breeders in the order best representing their population of interest or the latest findings in literature.

Reducing the relative importance of the adoption process predicted a higher proportion of dispersers as young wolves cannot be adopted by packs and therefore they remained floaters in the population (i.e., considered as “dispersers” in the model). The predicted populations had more resident individuals when the adoption process was more important. This process seems the one influencing the least our model outputs. However, we selected general model outputs to best represent the conservation state of the wolf populations, and this process may influence other elements of the wolf population that we missed. Predicting population with a reliable number of resident and dispersing individuals can be of high importance when modeling demographic processes where dispersers and floaters have a main role such as the colonization of new areas (e.g., Boyd et al., 1999, Pletsher et al., 1997).

One of the limitations of our IBM approach is that it is non-spatial. This greatly simplifies the use and the adaptation of the model to other populations as no animal-environment interactions are modeled and therefore no data regarding these interactions, which are sometimes hard to acquire, are needed. Parameters like equilibrium density, territory size, number of migrants and proportion of emigration that need to be defined by the user give one way to account for spatial constraints given by a particular environment, and these parameters can be changed to best represent the study area of interest. That said, we acknowledge that an explicit spatial mechanism would be very interesting to implement as wolf pack occupancy is mainly driven by exclusive territoriality (Cassidy et al., 2016), but the population division and affiliation into packs approximated spatiality in our model. To add more spatial constraints without changing the model structure, a new individual characteristic could be defined to represent distances between individuals based on their pack affiliation. Individuals from the same pack would be closer to each other than to other individuals, allowing a researcher to define short-distance dispersers vs long-distance dispersers (Louvrier et al., 2018), separate from the immigration/emigration process of our model. With a bit more work, the model could be turned into a spatially explicit IBM by including an explicit dispersal sub-model like that in Marucco and McIntire (2010) in place of the current dispersal sub-model. Model outputs were sensitive to only a few parameters, apart from those linked to the explored processes. Equilibrium density naturally affected the model predictions as this parameter represents, as stated above, one of the main spatial constraints influencing the simulated populations. This parameter triggered density-dependent events that occurred only when the landscape was fully occupied. Having more variability to trigger these events could likely reduce the influence of the equilibrium density parameter on model outputs and reduce the subjectivity of this trigger.

Building this IBM, we aimed to include all biological processes documented in the literature to best represent the wolf life cycle. Overall, we hope that our reproducible implementation of a modular and flexible IBM will contribute to the understanding, management and conservation of wolf populations by providing a scenario-testing and decision-making tool for ecologists and stakeholders, as well as a base model that can be adapted to simulate other canids and social species. The R language we used to code our IBM is largely used by ecologists and this should likely ease the model understanding and adaptation. For example, without much modification, our model could easily reproduce the life cycle of North American wolf populations with a few changes to the parameter values and the addition of a sub-model representing pack splitting (Jedrzejewski et al., 2004; Mech and Boitani, 2003). Users could also test hypotheses regarding assortative and disassortative mate choice for particular traits, relevant in hybridization studies (Fredrickson and Hedrick, 2006). Management actions (e.g., culling, sterilization) can as well be included in the model to test their effectiveness (Haight and Mech, 1997). The modular structure of our IBM allows the addition, removal or modification of only specific components of the model while keeping the other sub-models and the main structure the same. This model could be useful to other ecologists who could adapt it for their own specific research and management applications.

## Abbreviations

IBMs: individual-based models ID: identity
Fig.: Figure
ODD: Overview, Design concepts, and Details
popDensStd: wolf density per 1000 km^2^ standardized with mean and standard deviation from Cubaynes et al. (2014)
sd: standard deviation

## Acknowledgements

This study was supported by the French National Research Agency with a Grant ANR-16-CE02-0007 and by CNRS and the “Mission pour l’Interdisciplinarité” through the “Osez l’Interdisciplinarité” initiative. N. Santostasi was funded by a PhD grant from the Dept. of Environmental and Evolutionary Sciences of the University of Rome La Sapienza. O. Grente was funded by a PhD grant from the French Game and Wildlife Agency, and the French Office for Biodiversity. We thank Francesca Marucco and Eliot McIntire for letting us reuse some of their wolf IBM sub-models. We thank John Benson, Nolwenn Drouet-Hoguet and three anonymous reviewers for their comments on the manuscript.

## Appendix A

Complete description of the wolf individual-based model (IBM) following the Overview, Design concepts, and Details protocol (“ODD” protocol) developed by Grimm et al. (2006, 2010)

### Overview

#### Purpose

The wolf IBM aims to represent all non-spatial dynamics that happen in a wolf population, with a focus to detail pack dynamics, the change of status between disperser and residents, and the replacement of breeding individuals. The model also explores lesser-known processes of the wolf dynamics, namely: the pack dissolution following the loss of a breeder, the adoption of young dispersing individuals by packs, the establishment of new packs using the budding strategy, and the breeder replacement. These processes are known to happen in wolf populations but were rarely included in models (Marucco and McIntire 2010, Chapron et al. 2006; Pitt et al. 2003) due to the difficulty to parameterize or time them as little details are known on these processes. Other processes, better understood but also often over looked in models were also included: avoidance of inbreeding in packs where mating between wolves more related than two cousins is prevented as much as possible, density-dependent mortality for resident adults when the population is at equilibrium density, and movement of wolves in and out of the simulated population with possible immigrations and emigrations.

#### Entities, state variables, and scales

Entities in the model are wolf individuals. Each wolf has a unique ID, a sex (male or female), an age, a residence status (i.e., resident belonging to a pack or disperser), a pack ID if it belongs to a pack, a breeding status (i.e., breeding individual or not), a mother ID, a father ID, and a cohort ID (i.e., the year individuals are born in). ID, sex, mother ID, father ID and cohort ID never change during simulations. Age is updated each year. Residence status and pack ID change when an individual leaves or joins a pack. Breeding status changes when an individual becomes a breeder, either by replacing a missing one or by forming its own pack, or when an individual loses its breeder status after being replaced or after its pack broke apart. Packs are not considered entities in the model as most of the processes do not act on the whole pack at once (except pack dissolution). Packs are just a characteristic (via their ID) of the wolves. The model is non-spatial so the environment is not represented and wolves do not have a location. Temporal scale is a one-year time step.

#### Process overview and scheduling

In one year, all individuals go through the same series of sub-models (Fig. A1) and their state is modified according to the behavioral rules of each sub-model and their own characteristics. In order, these sub-models are: reproduction, aging, mortality, and change of resident/disperser and breeder/non-breeder status (Fig. A1). The change of the wolves’ residence and breeding status is represented with several sub-models that are: pack dissolution, replacement of breeding females by subordinates, dispersal, immigration/emigration, adoption, replacement of breeders by dispersers, establishment in pairs, establishment by budding, establishment alone, and replacement of breeding males by subordinates (Fig. A1). The order of the sub-models simulating breeder replacement is debated and therefore is explored through different model versions (M1, M2 and M3, Fig. A1). For simplicity, the order used to present the model in the ODD is the one of M1 (Fig. A1) but we do not state that this model version is more reliable than the other ones. In “reproduction,” new individuals (pups) are produced. In “aging,” the age of all individuals is updated. In “mortality,” different mortality probabilities affect different individuals based on their age, their residence status, and the number of existing packs relative to the equilibrium density. In “pack dissolution,” some packs dissolve based on the age composition in the pack, their number of individuals and of breeders. If some packs dissolve, the residence status of the individuals is updated to dispersers and they lose their pack ID. In “replacement of breeding females by subordinates,” female subordinates may replace the missing breeder in a pack and their breeding status is updated. In “dispersal,” packs with too many individuals force some individuals to leave the pack with different relative probabilities based on their age. These individuals have their residence status updated to dispersers and they lose their pack ID. In “immigration,” wolves from outside integrate into the population. In “emigration,” wolves from the simulated population leave the study area; similar to death, they are removed from the population. In “adoption,” young dispersing individuals may be adopted by small packs. These adoptees have their residence status updated to resident and they obtain the pack ID of their adopting pack. In “replacement of breeders by dispersers,” dispersing individuals may replace missing breeders in packs. These individuals have their residence status updated to resident, they obtain the pack ID they integrate and their breeding status is updated. In “establishment in pairs,” two dispersing individuals of the opposite sex establish themselves together to form a new pack. These individuals have their residence status updated to resident, they obtain a new and unique pack ID and their breeding status is updated. In “establishment by budding,” a dispersing individual and a subordinate from a pack establish a new pack. The former disperser has it residence status updated to resident, it obtains a new and unique pack ID and its breeding status is updated. The former subordinate obtains the same pack ID as its new partner and has its breeding status updated. In “establishment alone,” dispersers can establish themselves and form a pack alone. These individuals have their residence status updated to resident, they obtain a new and unique pack ID and their breeding status is updated. In “replacement of breeding males by subordinates,” male subordinates may replace the missing breeder in the pack so their breeding status is updated. At the end of the sub-model series, the information about the current population (i.e., the current characteristics of each individual) is saved and individuals go through the same loop of sub-models for as many years as simulated.

**Figure A1:**
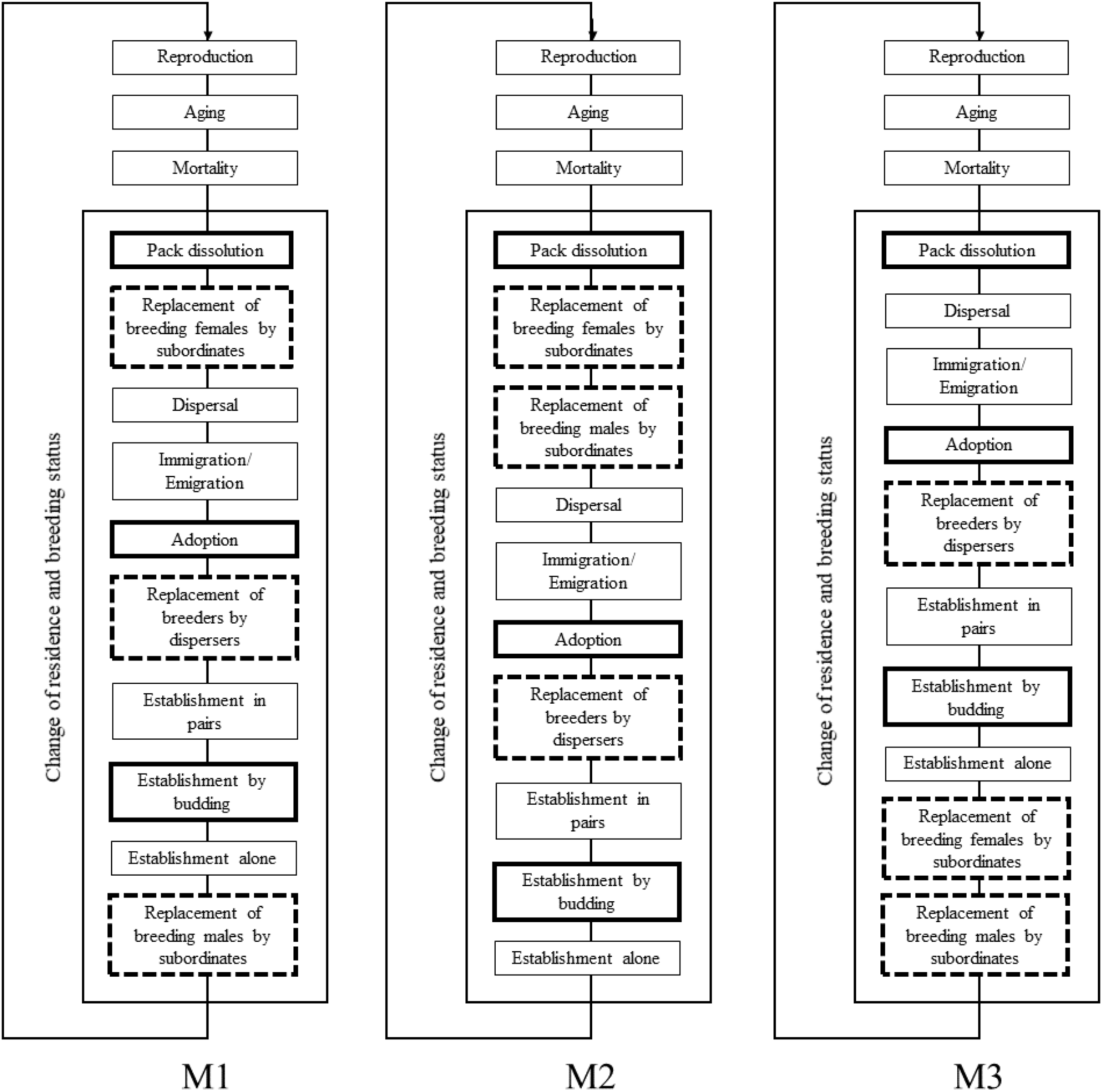
Diagram of three different versions of the wolf IBM (M1, M2 and M3). The sub-models in solid bold boxes are the lesser-known processes explored with different parameter values. The sub-models in dashed bold boxes are the lesser-known processes related to breeder replacement for which their order, instead of their parameter, is tested with the three versions of the sub-models series: M1, M2, and M3. When a wolf population is simulated with one of these model versions, individuals go through each sub-model of the loop all together, one sub-model at a time, for as long as the simulation lasts. The loop of sub-models represents a one-year time step.

### Design concepts

#### Basic principles

The life cycle of the wolf is represented through the reproduction, mortality, dispersal and establishment of the individuals, already defined in published wolf IBMs (Marucco and McIntire 2010, Chapron et al. 2016). However, extensive research in the literature has been done to understand, and then include in the IBM, all processes known to happen in wolf dynamics. Additionally to the fundamental processes, we included those related to the change of status between residents and dispersers, the access to the breeding status, the pairing between male and female breeders based on their relatedness and the residence and breeding status, the movement of wolves in and out of the population, and density-dependent processes. The model provides new details on the pack dynamics to mimic the gray wolf life cycle as best as possible. The model is non-spatial but the spatial distribution of individuals is represented through their pack affiliation. The life cycle represented in the IBM as well as the parameter values used are more adapted to wolf populations in central Europe (i.e., Alps) than for large North American populations (i.e. Canada, Alaska (USA)).

#### Emergence

Through reproduction and immigration, new individuals are added in the population. Individuals die and are removed from the population through different mortality and emigration processes. These changes in the population affect the total number of individuals. Processes affect wolves depending on their individual characteristics (i.e., age, sex, residence status, breeding status, etc.) and affect the distribution of the individuals in different classes (e.g., number of residents and dispersers, number of packs with two breeders in it).

#### Adaptation

Wolves live in packs and most of the processes coded in the model depend on the pack structure and the status of the individuals in the pack. The presence of zero, one of two breeders constrains the potential reproduction, pack dissolution, and replacement of breeding members. The total number of individuals in a pack constrains the pack dissolution, dispersal and adoption. The total number of established packs in the population also constrains some of the processes like the different probabilities of establishment (i.e., in pairs, by budding and alone) and one mortality process that have density dependent parameters.

#### Objectives

Wolves do not have an ultimate goal they need to fulfill over time. Individuals follow the behavioral rules of the different sub-models and respond to them according to their current characteristics and the current state of the packs.

#### Learning

There is no learning *per se* in the wolf IBM such as a learning of new skills (e.g., hunting prey taught by parents) but as individuals’ age and status change, the possibilities for the individual change. For example, wolves of age 1, 2 and 3 years old can be adopted but not at an older age. Only mature wolves (of age 2 and older) can become breeders and establish a new territory; pups of 1 year old cannot. Only mature breeding wolves can reproduce; mature subordinates cannot.

#### Prediction

Individuals know the current state of the population and individuals’ current characteristics but they cannot predict any future population or individual state nor any individuals’ actions.

#### Sensing

Wolves in packs have knowledge of all individual characteristics for the other members in their pack. Packs that can adopt young wolves can sense the presence of young dispersers. Dispersing individuals have access to the packs and their structure as replacement of missing breeders by dispersers and pairing with a subordinate from a pack is possible for these individuals. There is no sensing of the environment as there is no interaction with it.

#### Interaction

Wolves are social animals and therefore multiple interactions shape the life cycle of this species. Reproduction requires two breeding individuals from the same pack to produce pups. Pack dissolution and dispersal represent a loss of interactions between individuals that were members of a pack and become dispersers due to various factors. In the replacement of missing breeders, there is a choice among the mature subordinates of the pack or among mature dispersers that may be constrained by the presence of the other breeding wolf. During establishment in pairs or by budding, a disperser interacts with another disperser or a subordinate in a pack to create a new territory.

#### Stochasticity

Stochasticity is included in almost all components of the model. The number of pups produced per breeding pair, the number of individual dying, the maximum number of individual allowed in a pack, the number of immigrants arriving in the population and the number of emigrants leaving are generated using probabilities. The following processes also occur probabilistically: pack dissolution based on the number of breeding members remaining in the pack, adoption, dispersal according to the individuals’ age, and establishment by budding. Also, a density-dependent probability constrains the different types of establishment (i.e., in pairs, by budding and alone) as well as adult mortality when the population is at equilibrium density. The sex of the pups, the choice between unrelated individuals to replace missing breeders, the choice between young dispersers to be adopted, the choice between unrelated individuals to partner with a disperser to establish, and the choice of dispersing individuals that emigrate are done randomly. In the immigration process, as nothing is known about immigrating individuals, their sex and age (between a minimum and a maximum) is randomly chosen.

#### Collectives

Wolves belong to packs and their status of resident (i.e., inside a pack) or disperser (i.e., not belonging to a pack) affects almost all behavioral processes they follow. However, except for the pack dissolution, there is no process affecting the entire pack. As all individuals in the pack have different characteristics (i.e., age, sex, breeding status) they usually do not all respond in the same way.

#### Observation

The population is simulated for several years. Simulation outputs are available after each sub-model if needed or at the end of the whole series of sub-models at the end of the time step (i.e., at the end of the simulated year). The number of alive individuals with all their characteristics is available and many results can be extracted and derived from this population structure (e.g., the number of packs, the total abundance, the number of residents and dispersers, the number of breeders, the age distribution, etc.). We focused on outputs relevant for wolf conservation and management and defined six metrics. 1) At which year the population reached equilibrium density (i.e., in number of packs with a breeding pair). This output represents the speed of the population expansion and is a key element in areas that are being recolonized by the wolf. 2) The number of packs with two breeders. This metric is linked to the reproductive potential of the population and is of importance for management issues related to population growth. 3) The number of new packs formed in one year. This represents the pack turnover and the stability of the population that may affect hybridization and wolf-human conflicts. 4) The total number of individuals. 5) The proportion of residents and dispersers in the population. Population size is often required in management control and knowing the distribution of the resident/dispersing status of the individuals may help in understanding the population behavior. Finally, we looked at 6) the relatedness between the two breeders in each pack. Inbreeding avoidance plays a big part in the wolf life cycle, affecting the replacement of missing breeders and the creation of new packs. Often over looked because it is hard to simulate in non-individual-based models, this factor may indicate missing pieces in the models when not well represented.

### Details

#### Initialization

To launch the IBM, an initial wolf population is needed. We built a fictive population of 10 packs and 5 dispersing wolves, in a fictive environment that can hold 30 packs total (i.e., equilibrium density, Table A1). We created 5 packs with 2 breeders (one male and one female, 5 years old each) and 2 pups (one male and one female, 1 year old each); 3 packs with 2 breeders (one male and one female, 5 years old each), 1 yearling (one male, 2 years old) and 1 pup (one female, 1 year old); 2 packs with 2 breeders (one male and one female, 5 years old each) and 1 adult (one female, 3 years old); and 5 dispersers (3 females, 2 males, 2 years old each). This simple population was created for convenience but other initial populations can be easily defined by users. Table A1 lists all parameters and their values used in the model. These parameters can also be easily modified by the user. However, they represent the best data currently available in the literature for gray wolves in Europe, or elsewhere if not available for Europe. For lesser-known processes which parameters are explored (i.e., pack dissolution, adoption, establishment by budding), all tested values are listed (Table A1).

**Table A1:**
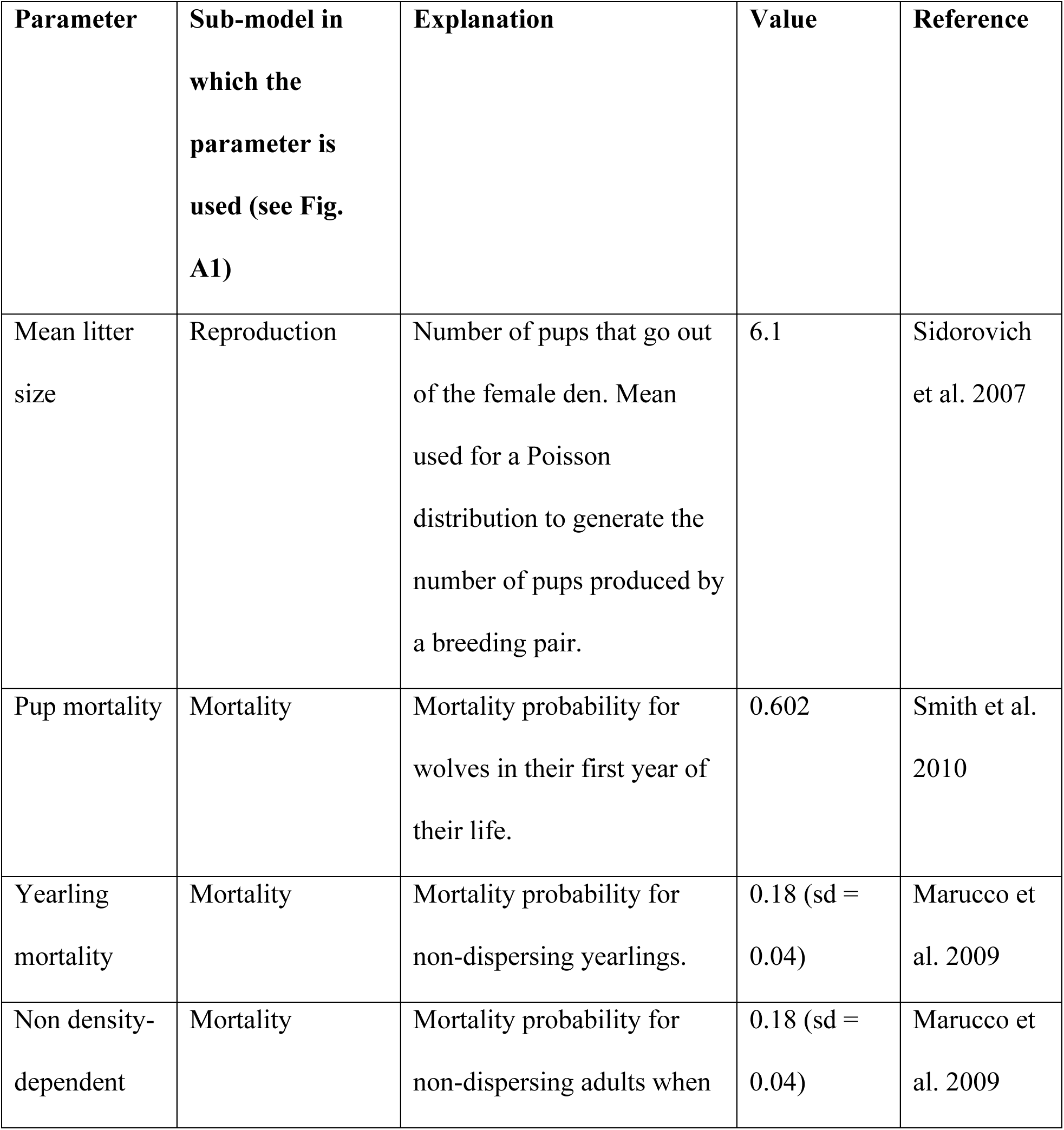
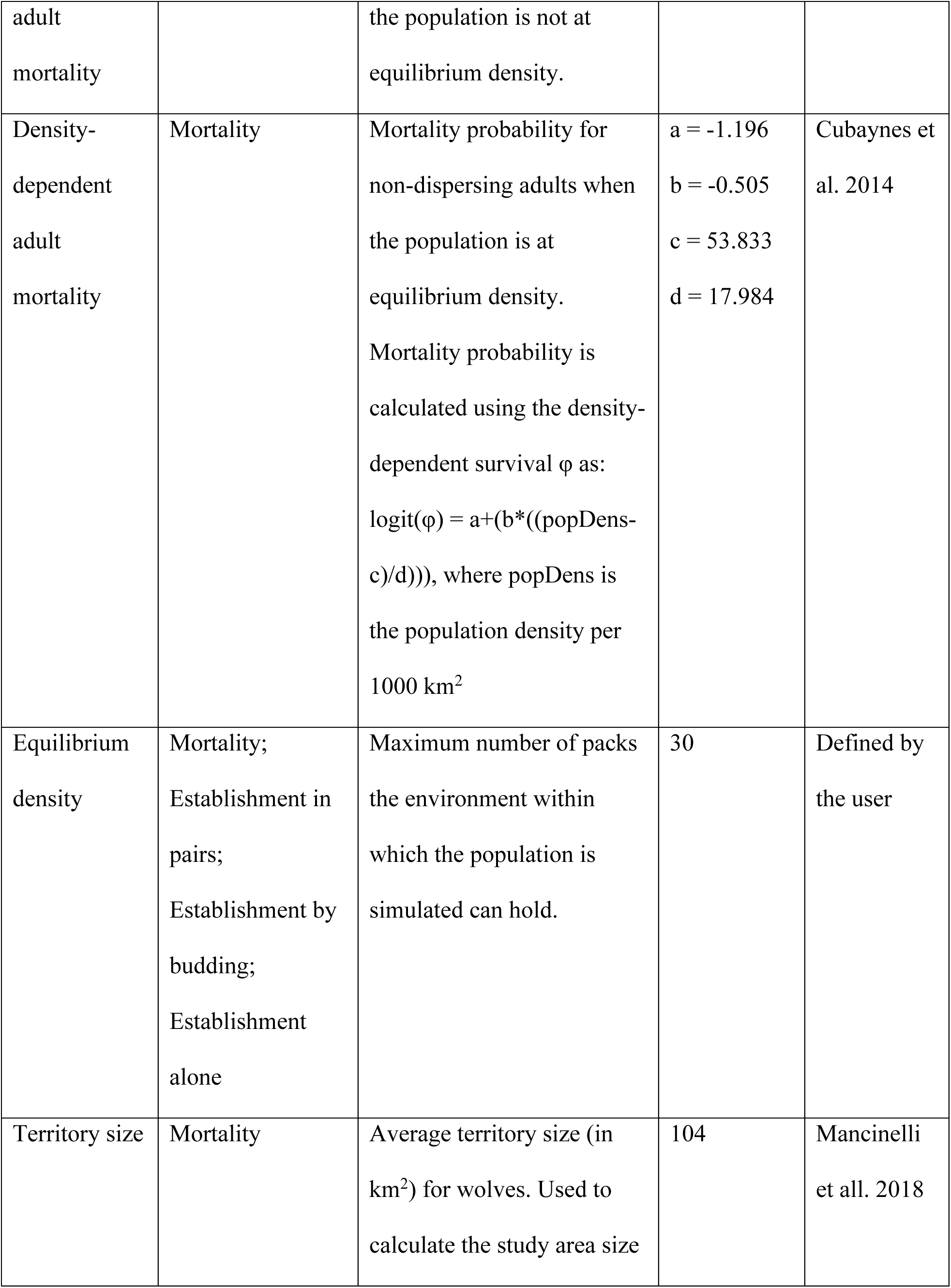
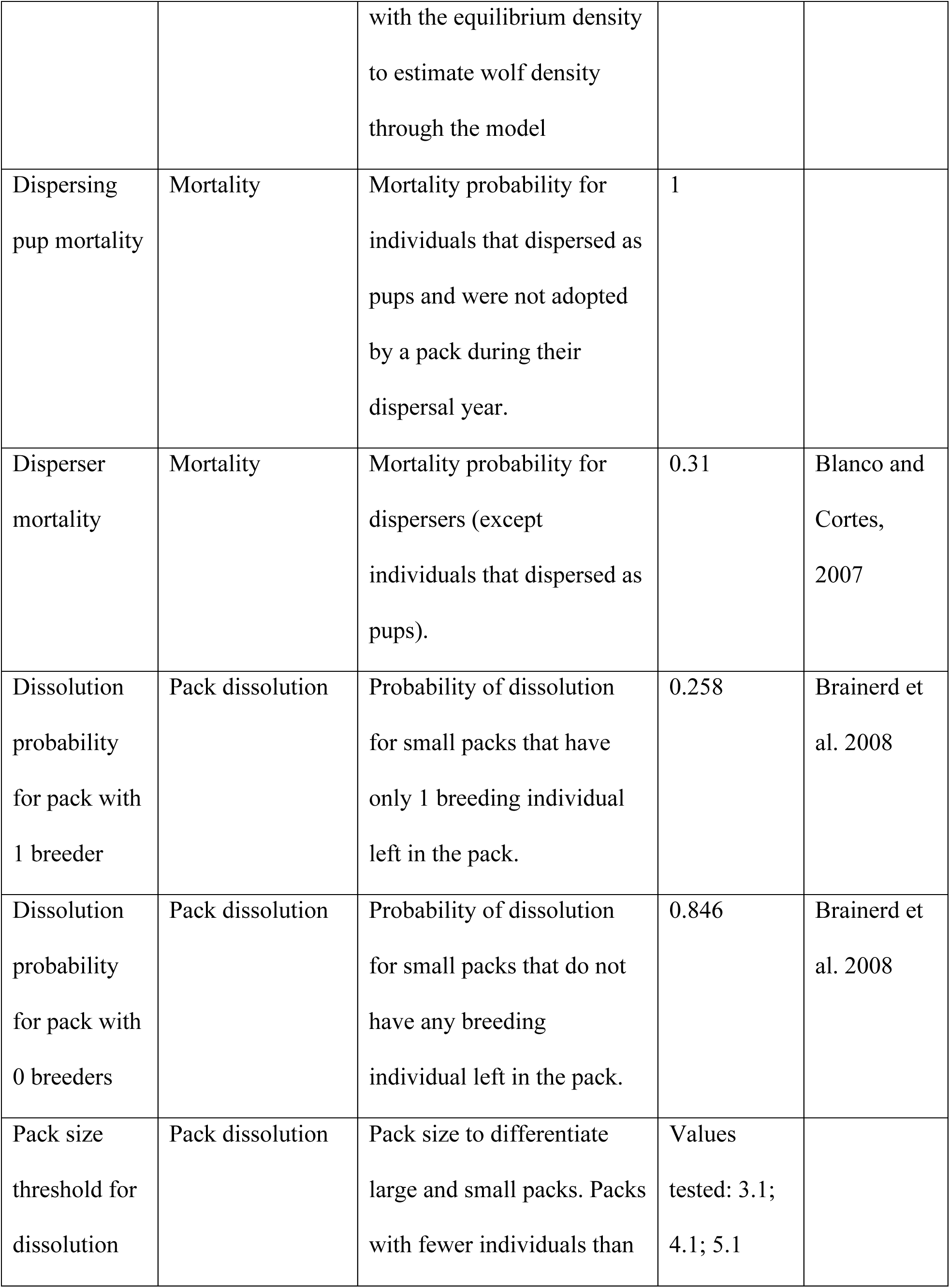
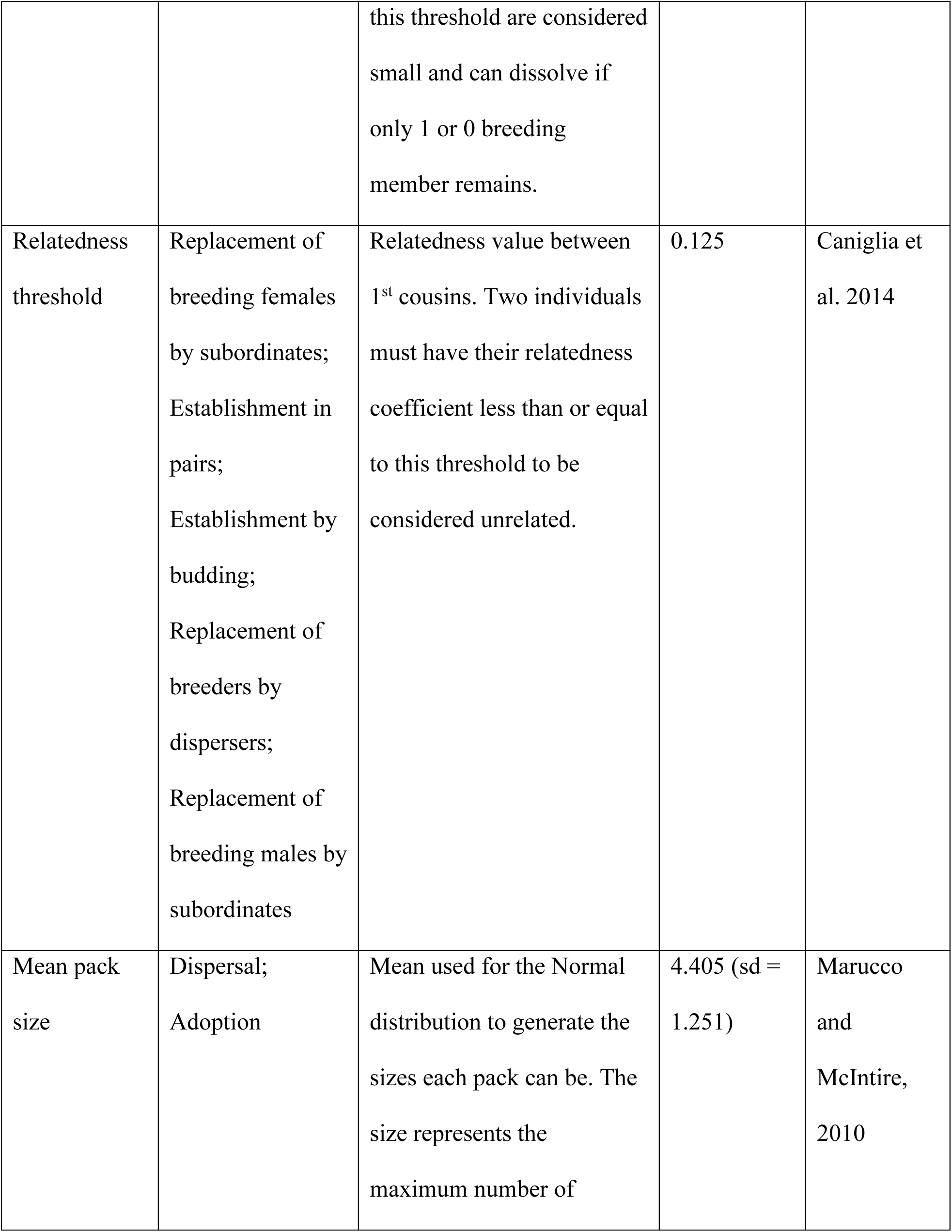
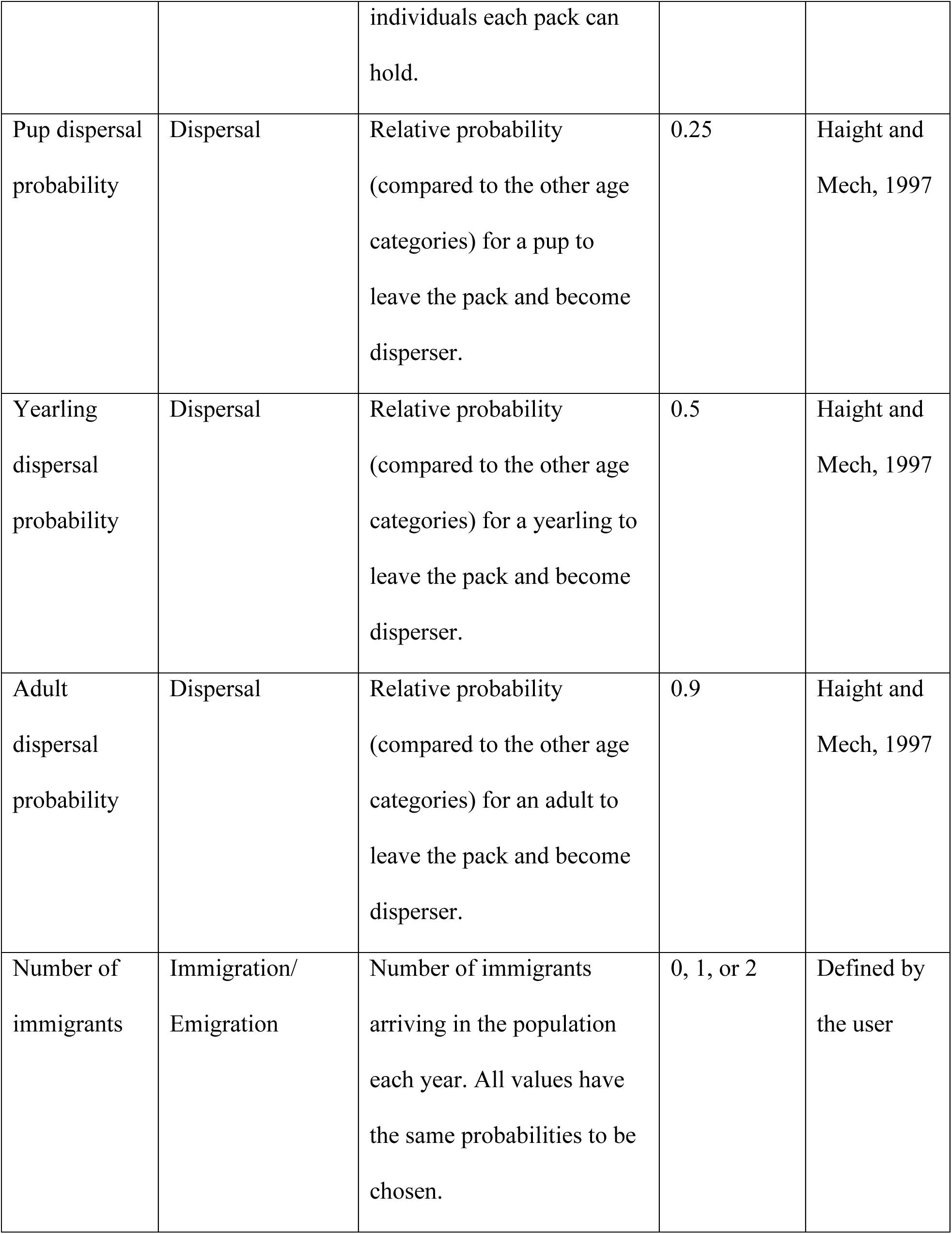
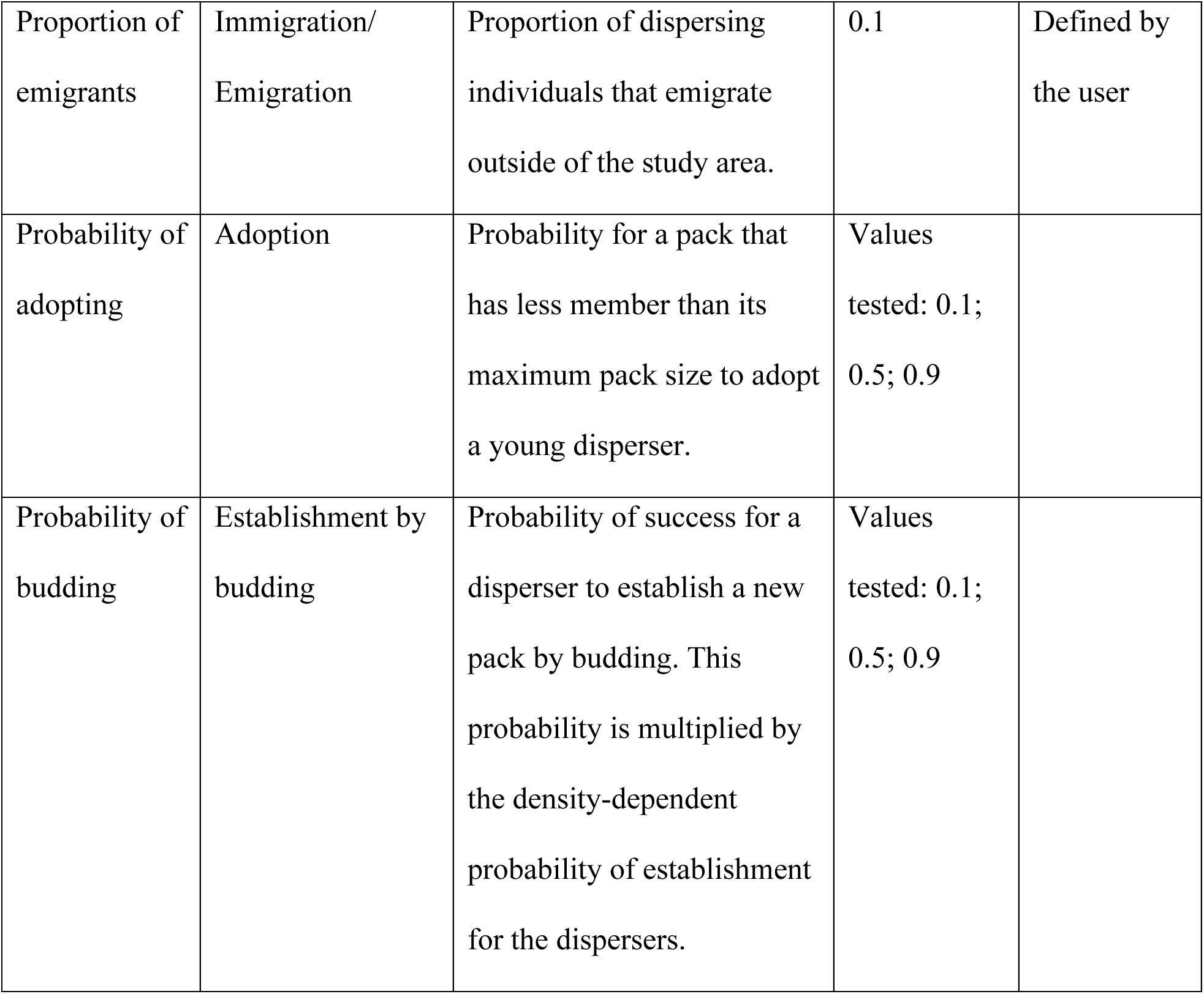
Parameters used in the wolf IBM. Probabilities and rates are estimates for a yearly time step.

#### Input data

There is no input data in the model. The environment is not represented and the initial population is not built using data.

#### Sub-models

*reproduction*: Every pack with both a breeding male and a breeding female reproduce (Marucco and McIntire 2010). The number of pups each pair has is drawn from a Poisson distribution (Chapron et al. 2016) with a mean of 6.1 (Sidorovich et al. 2007), representing the number of pups that emerge from the female den. Each pup receives a unique ID. The sex of each pup is randomly chosen as male or female with a 1:1 ratio (Sidorovich et al. 2007, Marucco and McIntire 2010). Their age is set to 0 as all individuals (including these newborn pups) will go through the “aging” sub-model next. Pups are considered residents, with the pack ID of their parents. They are not breeders. Their mother and father IDs are recorded and they obtain a cohort ID equal to the current simulated year (i.e., all pups born the same year have the same cohort ID). *aging*: All individuals age 1 year. Pups of the year are now 1 year old, yearlings are 2, and 3-year olds and older are adults. Individuals are considered mature at 2 years old (i.e., yearlings and adults).

*mortality*: There are 7 different mortality rates that take into account age, residence status and the total number of packs relative to equilibrium density. Mortality is applied individually to each wolf using a Bernoulli distribution which probability is sampled from a Normal distribution at each time step. Pups have a probability of 0.602 of dying (Smith et al. 2010). The mortality probability for non-dispersing yearlings is equal to 0.18 (sd = 0.04, Marucco et al. 2009). There are two types of mortality for non-dispersing adults that depend on the number of established packs in the population. If the number of established packs is below the number of packs at equilibrium density of the area, mortality is fixed and equal to that of the yearlings (i.e., 0.18 with sd = 0.04 (Marucco et al. 2009)). However, if the number of established packs is equal to the equilibrium density of the area, mortality is density-dependent. We used the equation linking wolf survival φ with wolf density from Cubaynes et al. (2014) to estimate the density-dependent mortality for non-dispersing adults: logit(φ) = 1.196 + (−0.505 * popDensStd), where popDensStd is the wolf density per 1000 km^2^ standardized with Cubaynes’ mean and standard deviation values (mean = 53.833, sd = 17.984). Wolf density is calculated as the total number of wolves, without considering the pups, divided by the area where the population is, estimated as the equilibrium density defined by the user multiplied by the wolf average territory size (104 km^2^, Mancinelli et al. 2018). Two mortality rates concern dispersing individuals. No pups can be dispersers and dispersing yearlings are individuals that dispersed the previous year (when they were pups due to a dissolution of their pack) but could not find a pack to adopt them during that year (otherwise they would be residents). We assumed these individuals are too young to survive by themselves and we defined a mortality probability equal to 1 to dispersing yearlings. All other dispersing wolves (i.e., adults) have a mortality probability equal to 0.31 (Blanco and Cortes 2007). We did not model senescence or any increase of the mortality probability with age. To represent a realistic age distribution in the population, the limit for wolves was the end of their 15^th^ year of simulation (Marucco and McIntire, 2010) and all individuals reaching 16 years old die.

*packDissolution*: Following the mortality event, packs whose social structure has been impacted by the loss of breeders may dissolve (Brainerd et al. 2008). Small packs with 1 breeding individual remaining will dissolve with a probability of 0.258 (Brainerd et al. 2008), and small packs with no breeder left dissolve with a probability of 0.846 (Brainerd et al. 2008). The pack size threshold to differentiate small and large packs was explored using the values 3.1, 4.1 and 5.1. In the specific case where both breeders died and only pups remain, the pack always dissolves as we assumed pups alone are unlikely to maintain a territory and so they disperse. When a pack dissolves, all former members of the pack become dispersers, they do not belong to a pack anymore and former breeding individuals lose their status.

*replaceBreedingFemBySub*: When a breeding female dies, she is most likely replaced by one of the female subordinates in her pack (most likely one of her daughters) (Caniglia et al. 2014; Jedrzejewski et al. 2005). When a pack is missing its breeding female, one of the mature females from the pack is randomly chosen to become breeder. Once the new breeding female is chosen we look at the relatedness between her and the current breeding male, if there is any. If there is a breeding male in the pack and he is closely related to the chosen female, he may be replaced (in sub-models *replaceBreederByDisp* and *replaceBreedingMalBySub*) by a disperser or a less related subordinate from the pack who will usurp the established breeding position (Mech and Boitani, 2003). The relatedness threshold chosen is the one of the first cousin (r = 0.125); a mating pair of breeding wolves can be no more closely related than cousins (e.g., no mating between siblings or parents and children) (Caniglia et al. 2014). This relatedness threshold is the same for all sub-models.

*dispersal*: When a pack has too many wolves, some are chased away and become dispersers. A maximum number of individuals is generated for each pack at each time step using a Normal distribution with a mean of 4.405 (sd = 1.251, Marucco and McIntire 2010). If the pack has more wolves than its maximum threshold, some individuals will leave the pack until the number of wolves in the pack is equal to its threshold. Breeding individuals cannot disperse. All the other wolves can disperse but with different relative probabilities based on their age. Pups may disperse with a relative probability of 0.25, yearlings may disperse with a relative probability of 0.5, and adults may disperse with a relative probability of 0.9 (Haight and Mech 1997). Wolves leaving the pack become dispersers and do not belong to the pack anymore.

*immigration*: Some wolves outside of the simulated population can arrive and interact with the other wolves. A user-determined number of immigrants will integrate with the population. The sex of the immigrants is randomly chosen (i.e., male or female with a 1:1 ratio). Their age is simulated using a truncated Poisson distribution of mean equal to 2 (with boundaries between 1 and 15) as yearlings are the most likely to disperse. Immigrants are dispersers, they do not belong to any pack yet and they are not breeders. As they were born outside the simulated population, they do not hold information about their mother, father or cohort IDs. Immigrant wolves will react the same way (i.e., follow the same sub-model rules) as the native wolves.

*emigration*: A proportion of the currently dispersing individuals, randomly chosen, leaves the simulated population via long-distance dispersal. These individuals will not come back and their disappearance is similar to simulating their death.

*adoption*: Packs which are not full (i.e., their number of individuals is below their maximum threshold) can adopt individuals. The probability with which these packs will adopt was explored using the values 0.1, 0.5, and 0.9. These packs can adopt individuals until they reach their maximum number of pack members. Only dispersers of 1, 2 and 3 years of age can be adopted by these packs. The order in which packs adopt dispersing individuals is random. Among potential adoptees, males are selected first. Then, if there are no more males and packs are still able to adopt, females are chosen. The choice among the males or among the females is random. Once young dispersers have been adopted, they become residents and belong to the pack that adopted them.

*replaceBreederByDisp*: Missing breeders in packs can be replaced by dispersers. First, we look at the packs missing breeding females. Mature female dispersers can become breeding females. If there is already a breeding male in the pack, we exclude the dispersing females that are closely related to the breeding male from the potential successors. Then, a female is randomly chosen among the unrelated ones to integrate into the pack. All selected females become residents and breeders of their assigned pack, and belong to the pack they joined. The order in which packs fill breeding female positions is random. Next, the same process is used to replace missing male breeders with mature male dispersers. If there are packs where the missing breeding female was replaced by a subordinate (in *replaceBreedingFemBySub*) during the time step and the current breeding male was too related to her, an unrelated, mature male disperser may usurp the established male breeder (Mech and Boitani, 2003). The breeding males replaced by dispersers are dismissed from their position and become subordinates in their own pack.

*establishPairs*: A male disperser and a female disperser can establish a new pack together if they are mature and not closely related. In addition, this is only possible if the number of existing packs is not already equal to equilibrium density. If the area is not already full, there is a density-dependent probability for dispersers to establish in pairs defined by a Bernoulli distribution with a probability equal to the number of packs that can be created until reaching equilibrium density divided by the equilibrium density. The more packs there are, the less likely it is that two dispersers establish themselves in pairs. Once a male and a female disperser have established a new pack, they both become breeders and residents, and obtain the same, new and unique pack ID. The order for the choice of males and females among the available mature dispersers is random.

*establishBudding*: Budding is when a disperser and a mature subordinate wolf from an existing pack establish a new pack together. Like establishment in pairs, budding is possible only if the number of packs has not reached equilibrium density. The probability for a disperser to bud is the density-dependent probability for establishment in pairs multiplied by a probability of budding. We explored this last probability and tested values equal to 0.1, 0.5 and 0.9. Only mature dispersers can bud, and only with a non-breeding, mature resident of the opposite sex that is not closely related. Once a disperser and a subordinate wolf bud, they both become breeders and residents, and obtain the same, new and unique pack ID. The order for the choice of males and females among the available mature dispersers and subordinates in packs is random. *establishAlone*: If the area is not at equilibrium density, remaining mature dispersers that could not establish themselves in pairs or by budding can establish themselves alone. The probability of this is also density-dependent, and is the same as the probability of the establishment in pairs. Once they create their own pack, wolves become breeders and residents, and obtain a new and unique pack ID.

*replaceBreedingMalBySub*: When a breeding male is missing, one of the mature, male subordinates in the pack can take over. If there are several subordinates that are eligible to become successors, the male least related to the current breeding female is chosen. If there are several subordinate males that are the least related, or if there is no breeding female, one is selected randomly. If the breeding female is too related to the newly chosen breeding male, the mature, female subordinate who is least related to the new breeding male can usurp the current breeding female and the current breeding female is dismissed (i.e., becomes subordinate). If there are several mature female subordinates that are the least related, one is selected randomly. In the particular case where there was a missing breeding female who was replaced by a subordinate (in *replaceBreedingFemBySub*) during the time step and she was too related to the current breeding male, one of the less related male subordinates can take over the male breeding position. All of these rules mimic the fact that wolves change partners to avoid inbreeding, except when there is no other choice (Mech and Boitani, 2003). Once new breeding individuals are chosen, they will be able to mate the next year.

## Appendix B

All 81 model scenarios tested by combining the 3 parameterizations of the pack dissolution process (Table 2, main text), with the 3 parameterizations of the adoption process (Table 2, main text), with the 3 parameterizations of the establishment by budding process (Table 2, main text), and with the 3 model versions for the breeder replacement process (Fig. 1, main text).

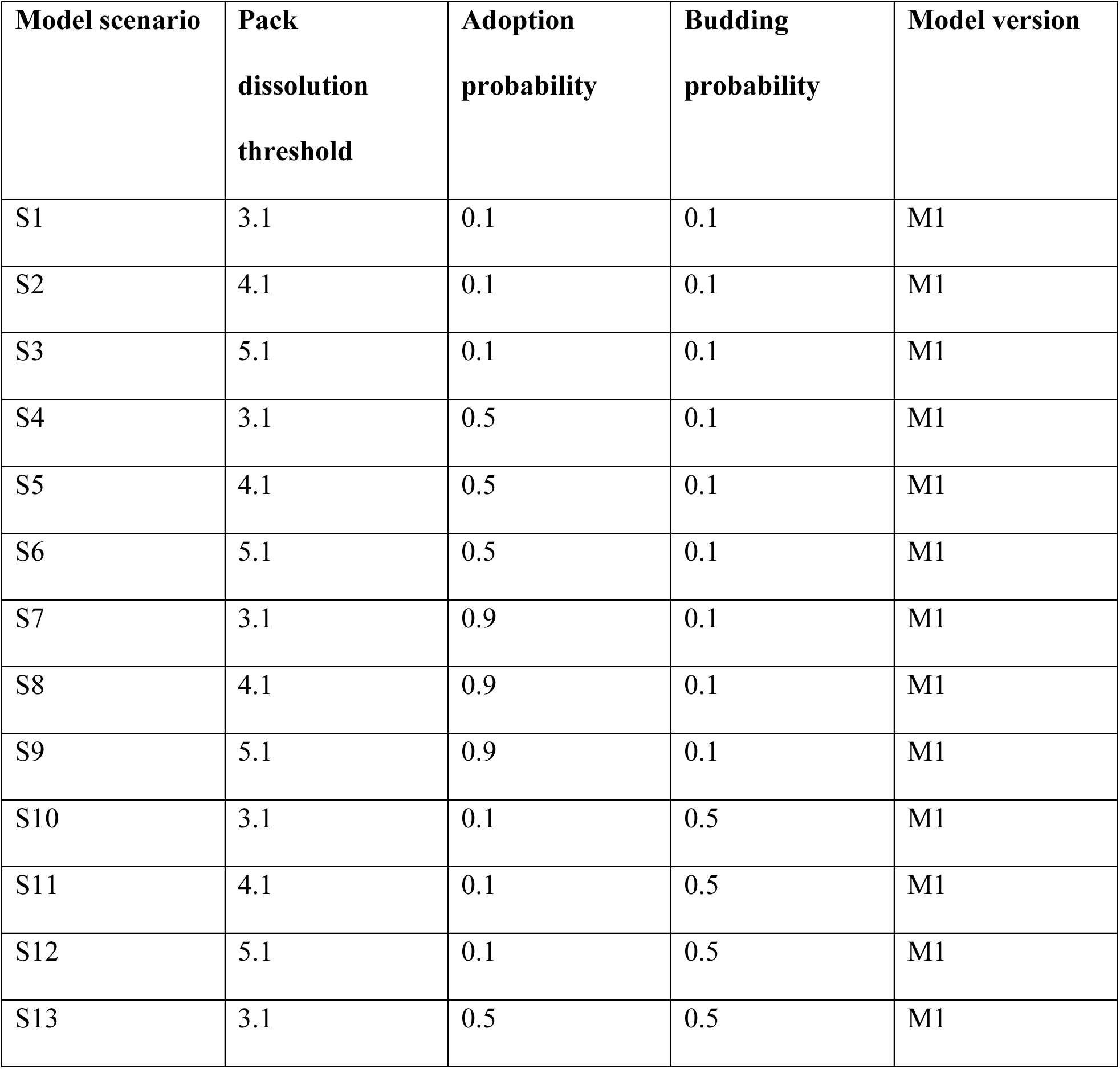
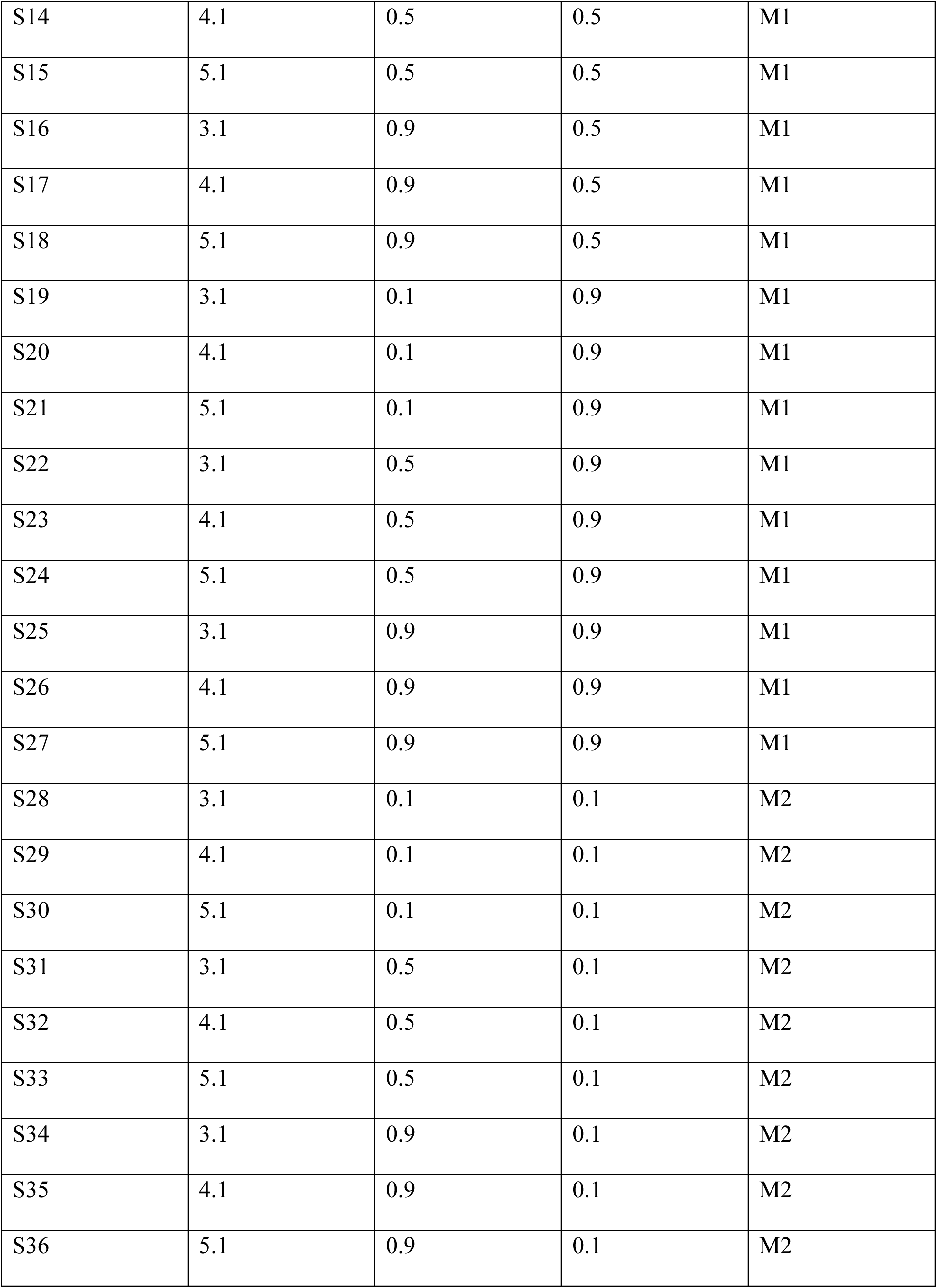
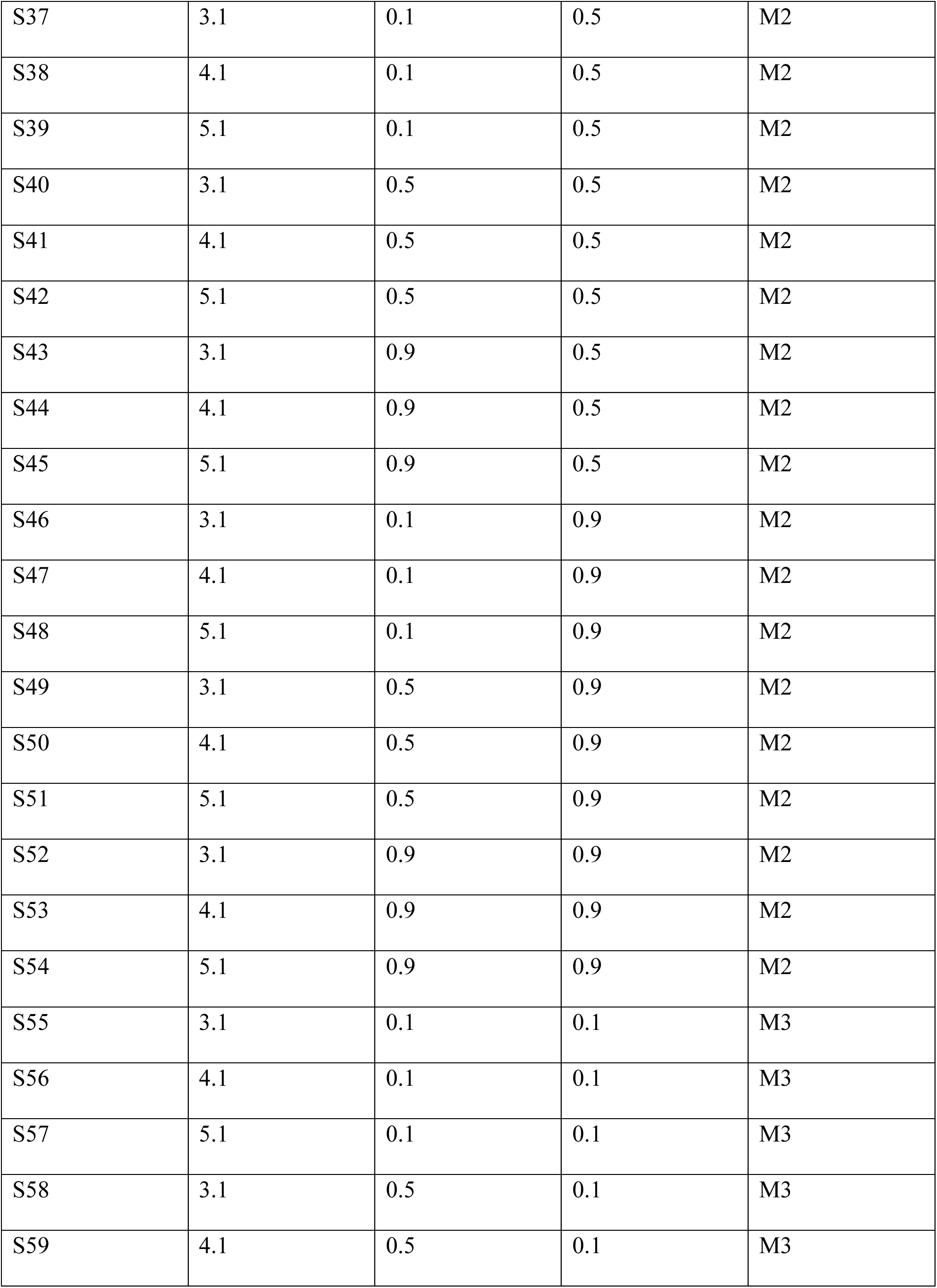
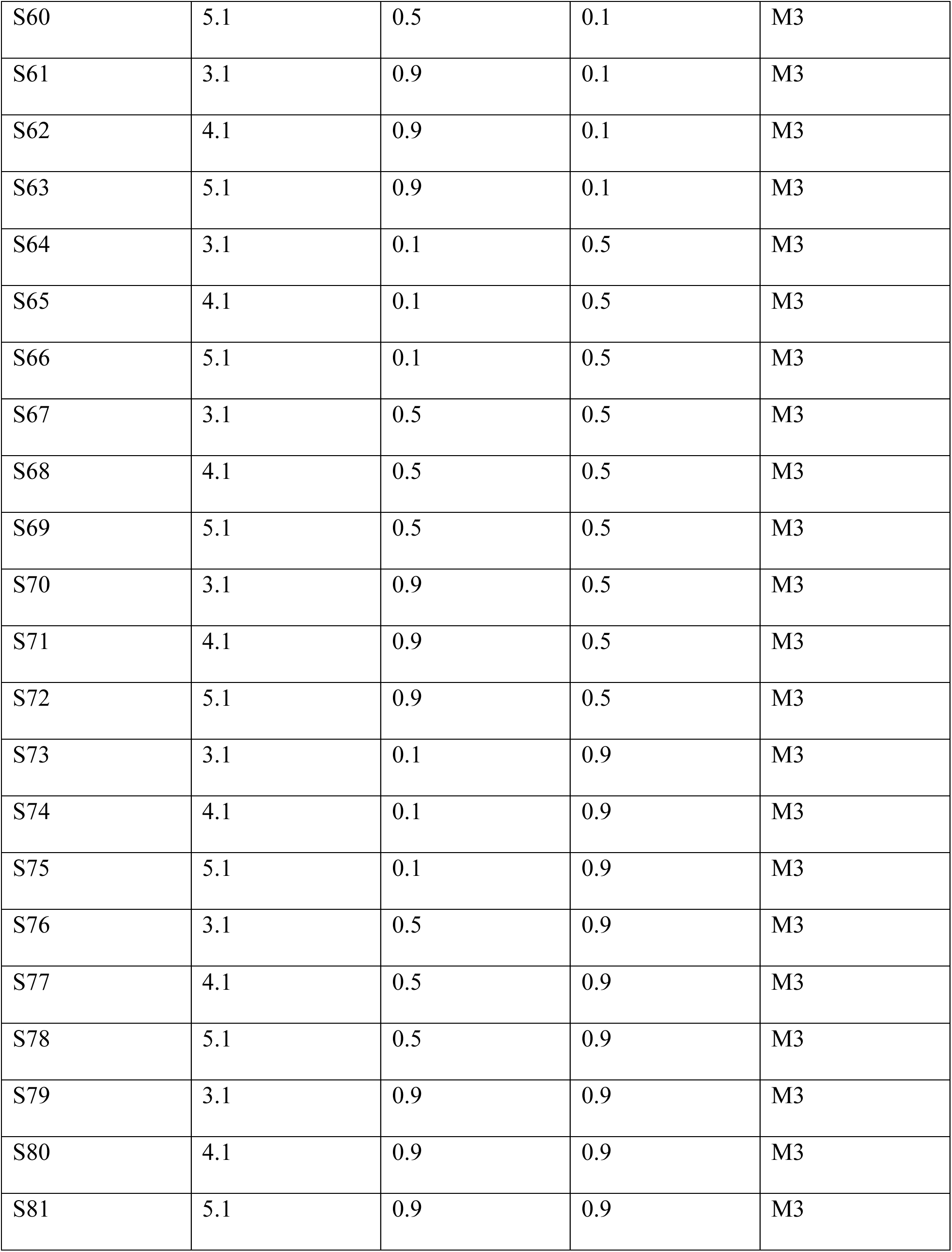

| **Model scenario** | **Pack dissolution threshold** | **Adoption probability** | **Budding probability** | **Model version** |
| --- | --- | --- | --- | --- |
| S1 | 3.1 | 0.1 | 0.1 | M1 |
| S2 | 4.1 | 0.1 | 0.1 | M1 |
| S3 | 5.1 | 0.1 | 0.1 | M1 |
| S4 | 3.1 | 0.5 | 0.1 | M1 |
| S5 | 4.1 | 0.5 | 0.1 | M1 |
| S6 | 5.1 | 0.5 | 0.1 | M1 |
| S7 | 3.1 | 0.9 | 0.1 | M1 |
| S8 | 4.1 | 0.9 | 0.1 | M1 |
| S9 | 5.1 | 0.9 | 0.1 | M1 |
| S10 | 3.1 | 0.1 | 0.5 | M1 |
| S11 | 4.1 | 0.1 | 0.5 | M1 |
| S12 | 5.1 | 0.1 | 0.5 | M1 |
| S13 | 3.1 | 0.5 | 0.5 | M1 |
| S14 | 4.1 | 0.5 | 0.5 | M1 |
| S15 | 5.1 | 0.5 | 0.5 | M1 |
| S16 | 3.1 | 0.9 | 0.5 | M1 |
| S17 | 4.1 | 0.9 | 0.5 | M1 |
| S18 | 5.1 | 0.9 | 0.5 | M1 |
| S19 | 3.1 | 0.1 | 0.9 | M1 |
| S20 | 4.1 | 0.1 | 0.9 | M1 |
| S21 | 5.1 | 0.1 | 0.9 | M1 |
| S22 | 3.1 | 0.5 | 0.9 | M1 |
| S23 | 4.1 | 0.5 | 0.9 | M1 |
| S24 | 5.1 | 0.5 | 0.9 | M1 |
| S25 | 3.1 | 0.9 | 0.9 | M1 |
| S26 | 4.1 | 0.9 | 0.9 | M1 |
| S27 | 5.1 | 0.9 | 0.9 | M1 |
| S28 | 3.1 | 0.1 | 0.1 | M2 |
| S29 | 4.1 | 0.1 | 0.1 | M2 |
| S30 | 5.1 | 0.1 | 0.1 | M2 |
| S31 | 3.1 | 0.5 | 0.1 | M2 |
| S32 | 4.1 | 0.5 | 0.1 | M2 |
| S33 | 5.1 | 0.5 | 0.1 | M2 |
| S34 | 3.1 | 0.9 | 0.1 | M2 |
| S35 | 4.1 | 0.9 | 0.1 | M2 |
| S36 | 5.1 | 0.9 | 0.1 | M2 |
| S37 | 3.1 | 0.1 | 0.5 | M2 |
| S38 | 4.1 | 0.1 | 0.5 | M2 |
| S39 | 5.1 | 0.1 | 0.5 | M2 |
| S40 | 3.1 | 0.5 | 0.5 | M2 |
| S41 | 4.1 | 0.5 | 0.5 | M2 |
| S42 | 5.1 | 0.5 | 0.5 | M2 |
| S43 | 3.1 | 0.9 | 0.5 | M2 |
| S44 | 4.1 | 0.9 | 0.5 | M2 |
| S45 | 5.1 | 0.9 | 0.5 | M2 |
| S46 | 3.1 | 0.1 | 0.9 | M2 |
| S47 | 4.1 | 0.1 | 0.9 | M2 |
| S48 | 5.1 | 0.1 | 0.9 | M2 |
| S49 | 3.1 | 0.5 | 0.9 | M2 |
| S50 | 4.1 | 0.5 | 0.9 | M2 |
| S51 | 5.1 | 0.5 | 0.9 | M2 |
| S52 | 3.1 | 0.9 | 0.9 | M2 |
| S53 | 4.1 | 0.9 | 0.9 | M2 |
| S54 | 5.1 | 0.9 | 0.9 | M2 |
| S55 | 3.1 | 0.1 | 0.1 | M3 |
| S56 | 4.1 | 0.1 | 0.1 | M3 |
| S57 | 5.1 | 0.1 | 0.1 | M3 |
| S58 | 3.1 | 0.5 | 0.1 | M3 |
| S59 | 4.1 | 0.5 | 0.1 | M3 |
| S60 | 5.1 | 0.5 | 0.1 | M3 |
| S61 | 3.1 | 0.9 | 0.1 | M3 |
| S62 | 4.1 | 0.9 | 0.1 | M3 |
| S63 | 5.1 | 0.9 | 0.1 | M3 |
| S64 | 3.1 | 0.1 | 0.5 | M3 |
| S65 | 4.1 | 0.1 | 0.5 | M3 |
| S66 | 5.1 | 0.1 | 0.5 | M3 |
| S67 | 3.1 | 0.5 | 0.5 | M3 |
| S68 | 4.1 | 0.5 | 0.5 | M3 |
| S69 | 5.1 | 0.5 | 0.5 | M3 |
| S70 | 3.1 | 0.9 | 0.5 | M3 |
| S71 | 4.1 | 0.9 | 0.5 | M3 |
| S72 | 5.1 | 0.9 | 0.5 | M3 |
| S73 | 3.1 | 0.1 | 0.9 | M3 |
| S74 | 4.1 | 0.1 | 0.9 | M3 |
| S75 | 5.1 | 0.1 | 0.9 | M3 |
| S76 | 3.1 | 0.5 | 0.9 | M3 |
| S77 | 4.1 | 0.5 | 0.9 | M3 |
| S78 | 5.1 | 0.5 | 0.9 | M3 |
| S79 | 3.1 | 0.9 | 0.9 | M3 |
| S80 | 4.1 | 0.9 | 0.9 | M3 |
| S81 | 5.1 | 0.9 | 0.9 | M3 |

## Appendix C

All model outputs from the 81 model scenarios tested (see Appendix B). Output shown over time (i.e., line figures) present the mean values and their 95 % confidence intervals, per year, from the 200 replicates for each model scenario. Boxplots present model outputs extracted at the end of the last simulated year from all 200 replicates for each model scenario tested. Lines and boxplots are color-coded (and ranked for boxplots only) according to the different values or model versions tested to explore the sub-models simulating lesser-known wolf dynamics processes: a) pack dissolution (see Table 2, main text), b) adoption (see Table 2, main text), c) establishment by budding (see Table 2, main text), and d) breeder replacement (see Fig. 1, main text).

**Figure C1:**
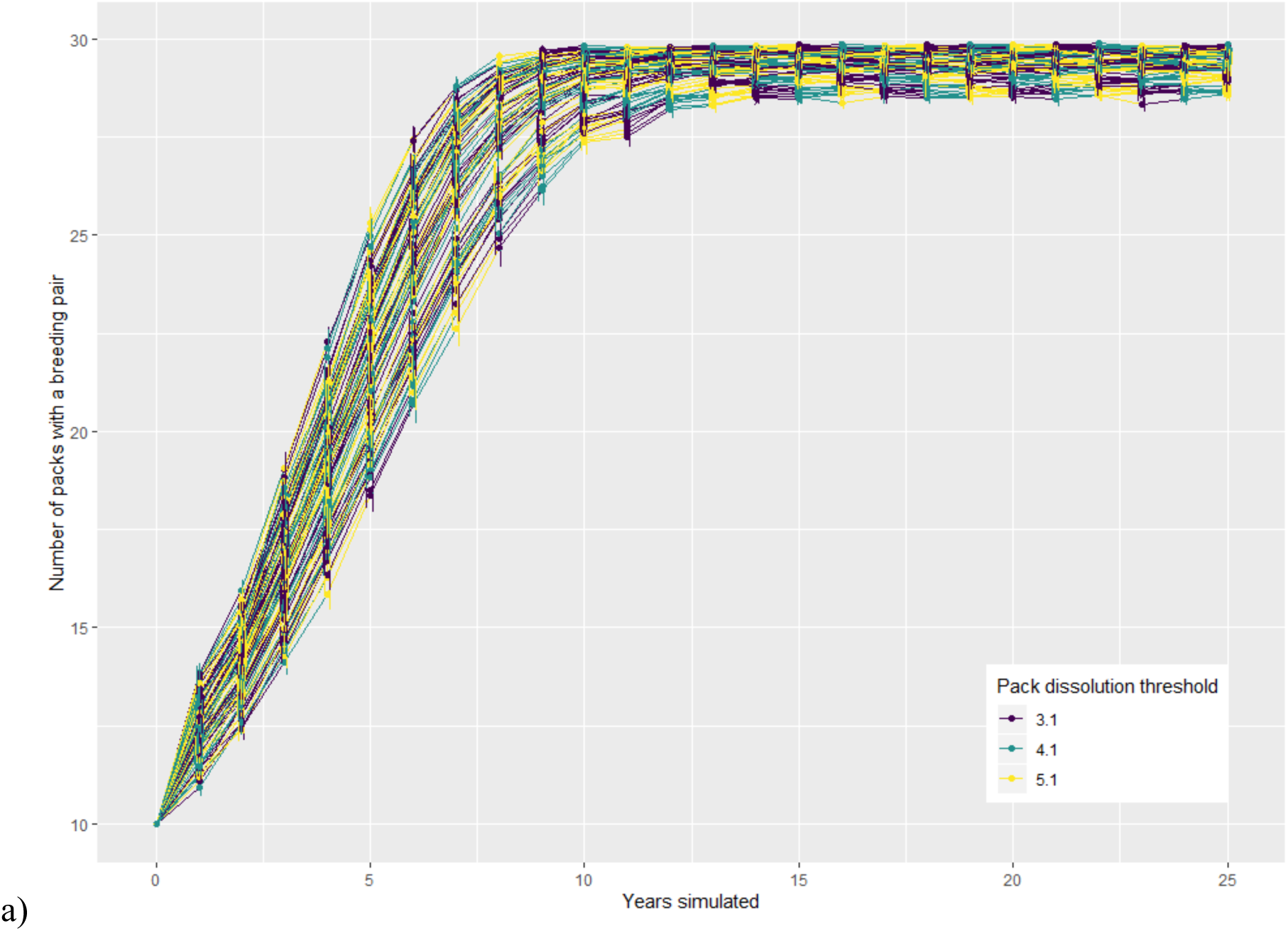
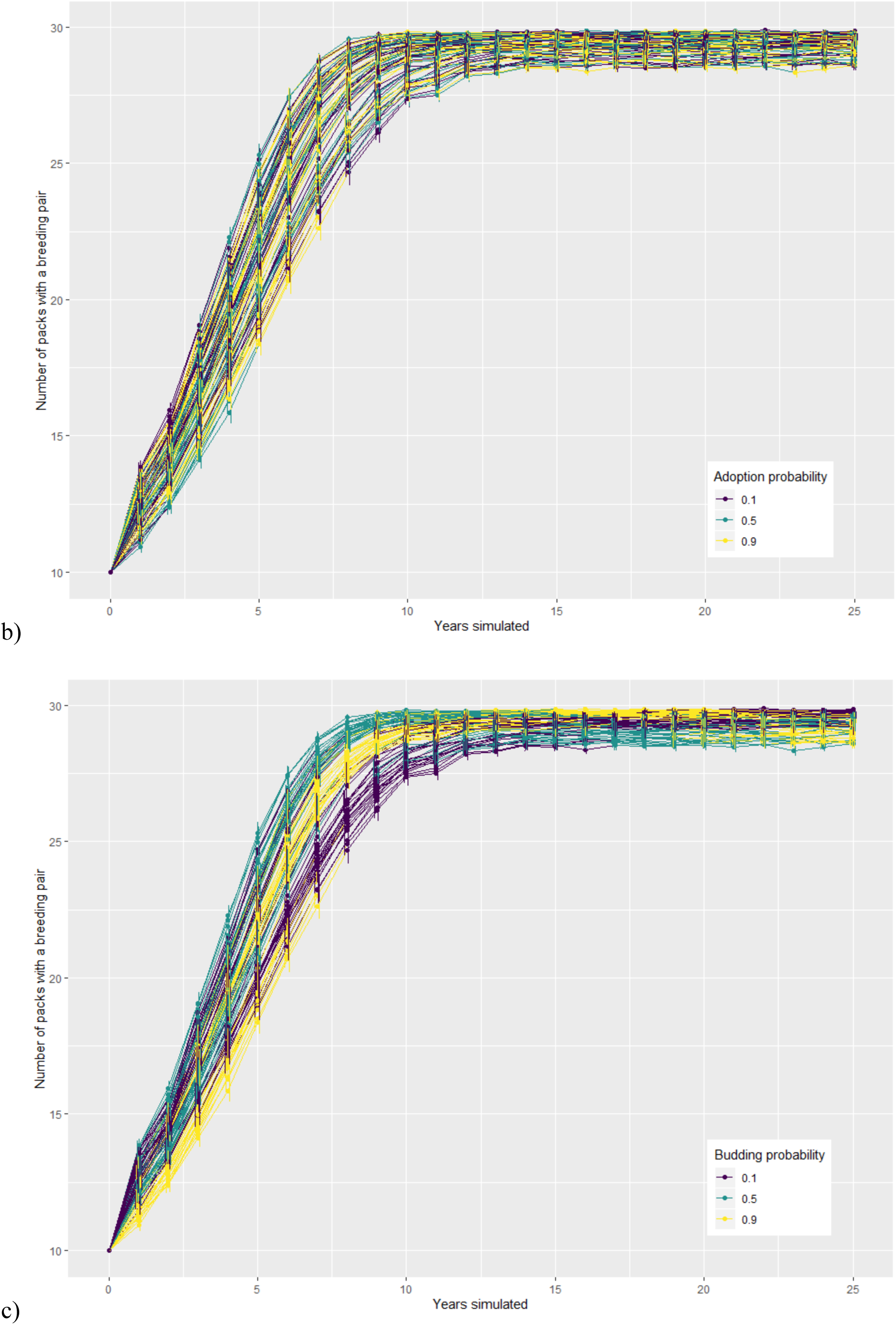
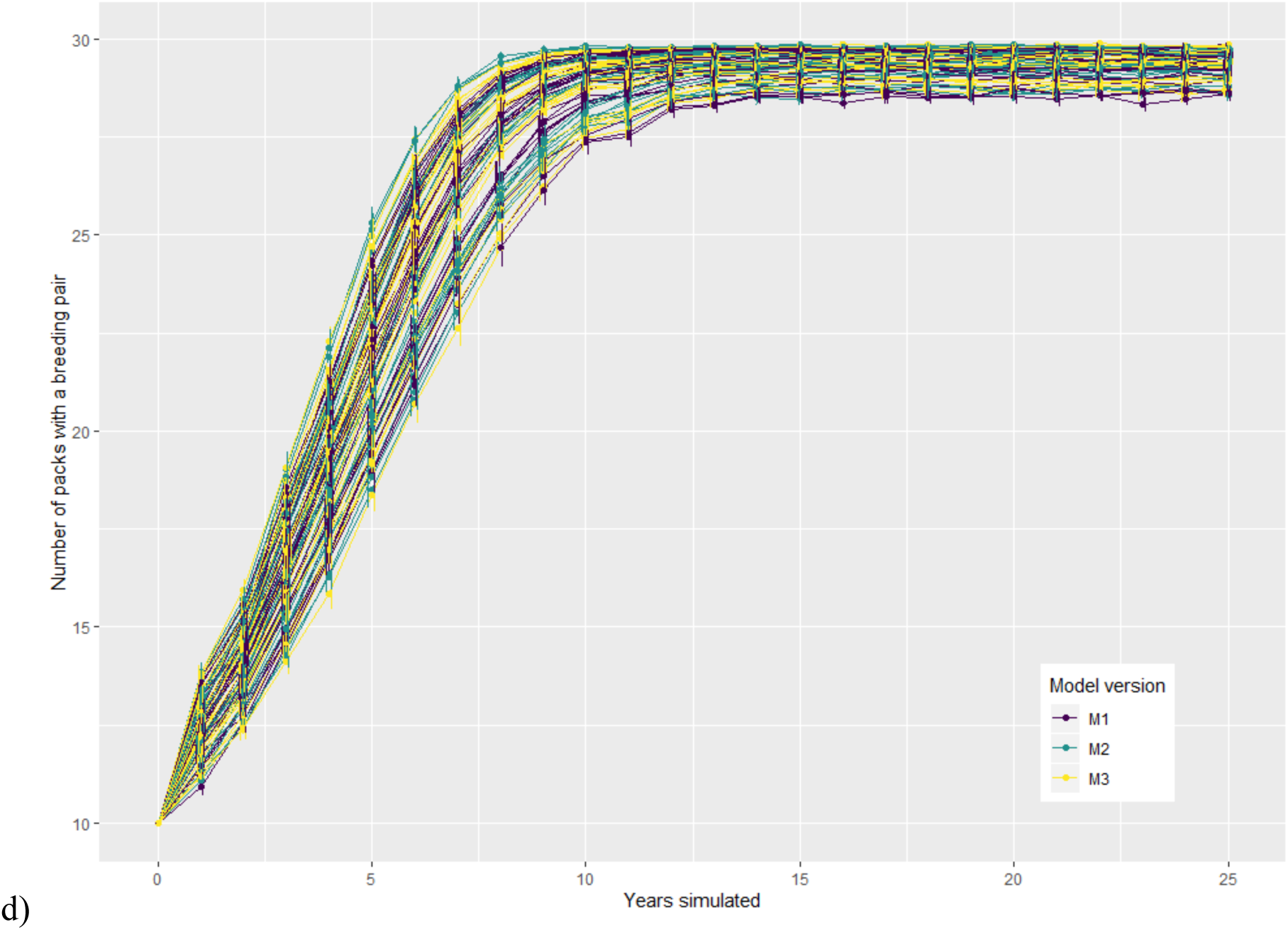
Number of packs with a breeding pair over the simulated years for each model scenario.

**Figure C2:**
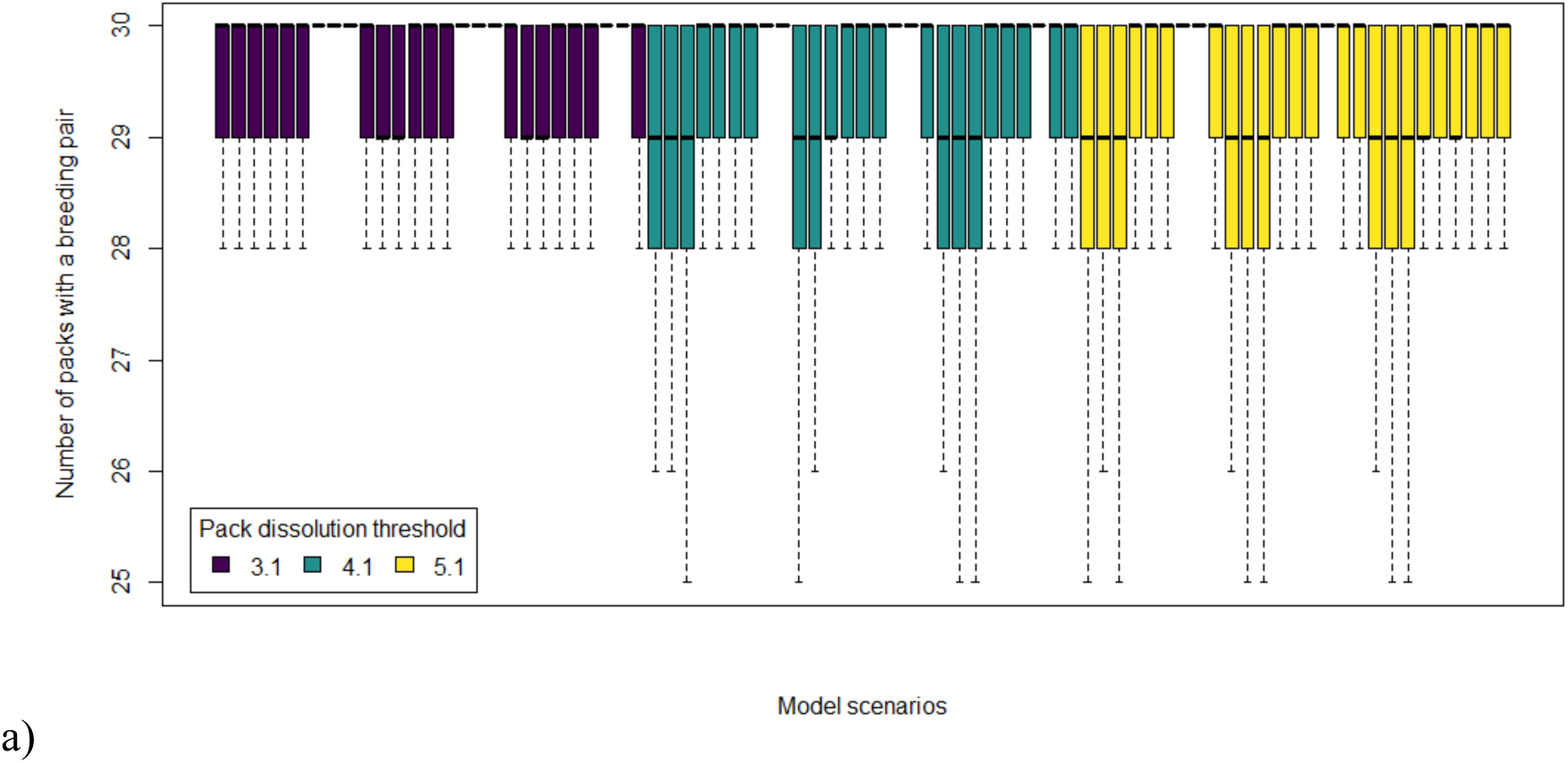
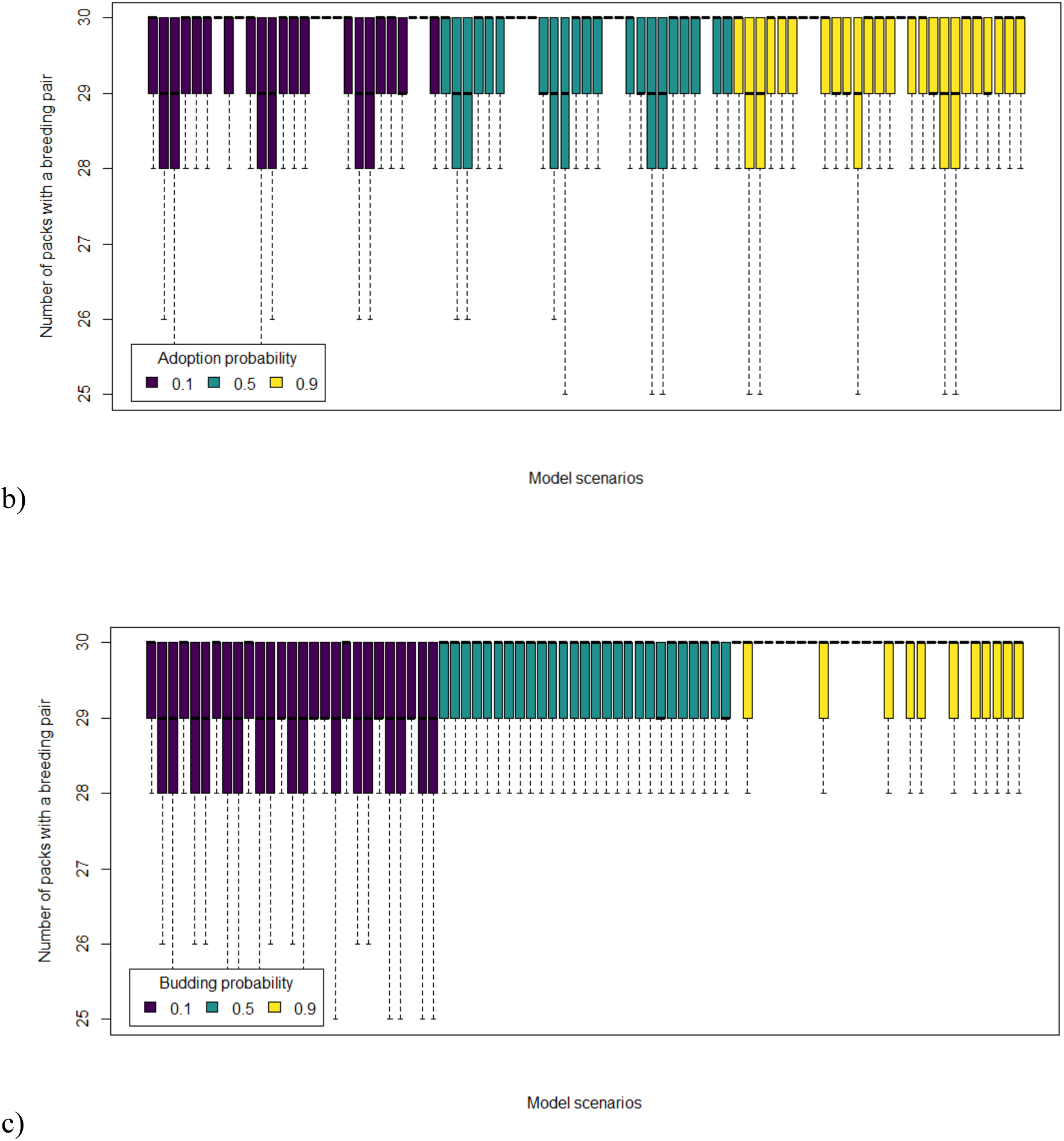
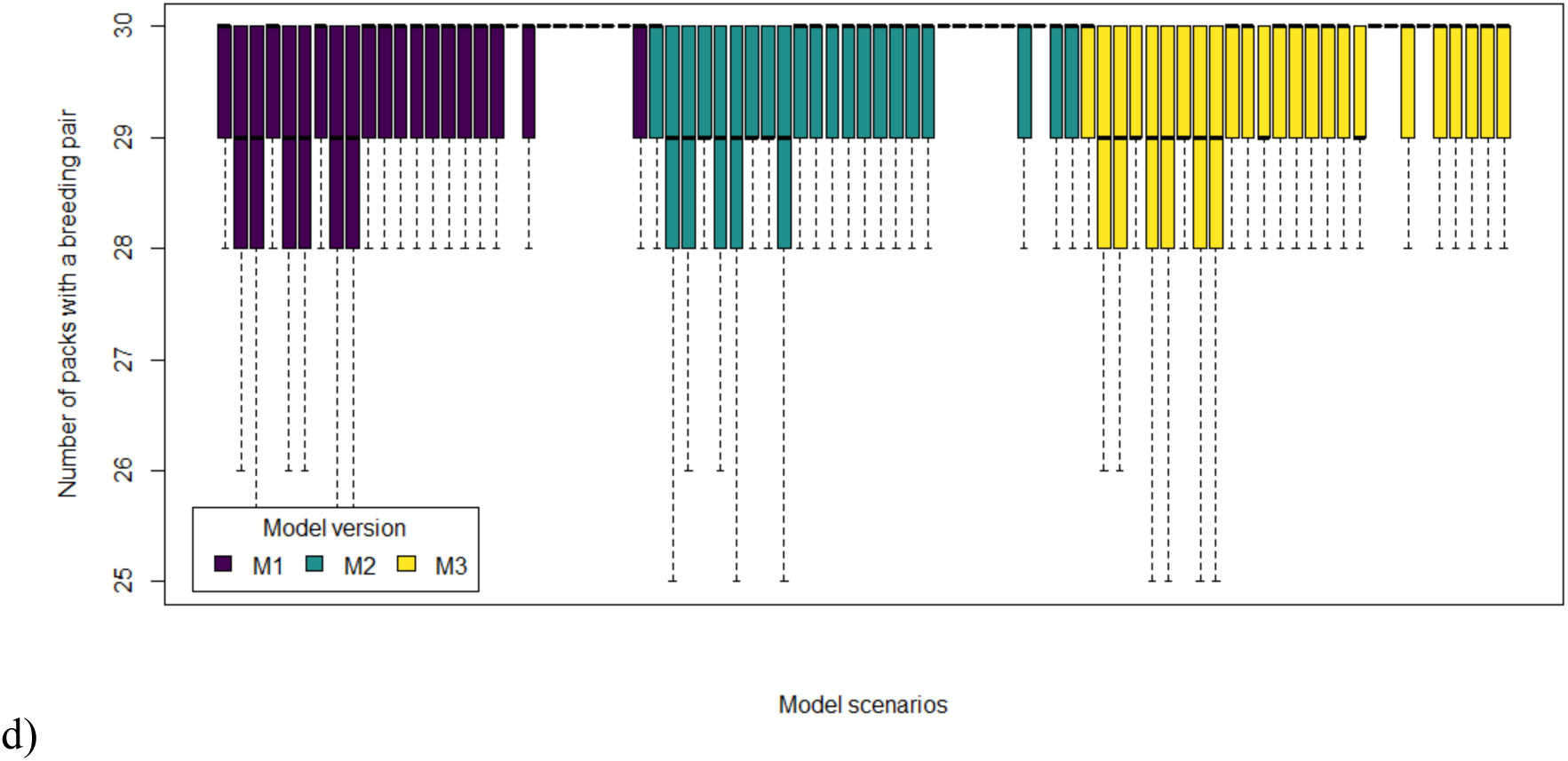
Number of packs with a breeding pair at the end of last simulated year for each model scenario.

**Figure C3:**
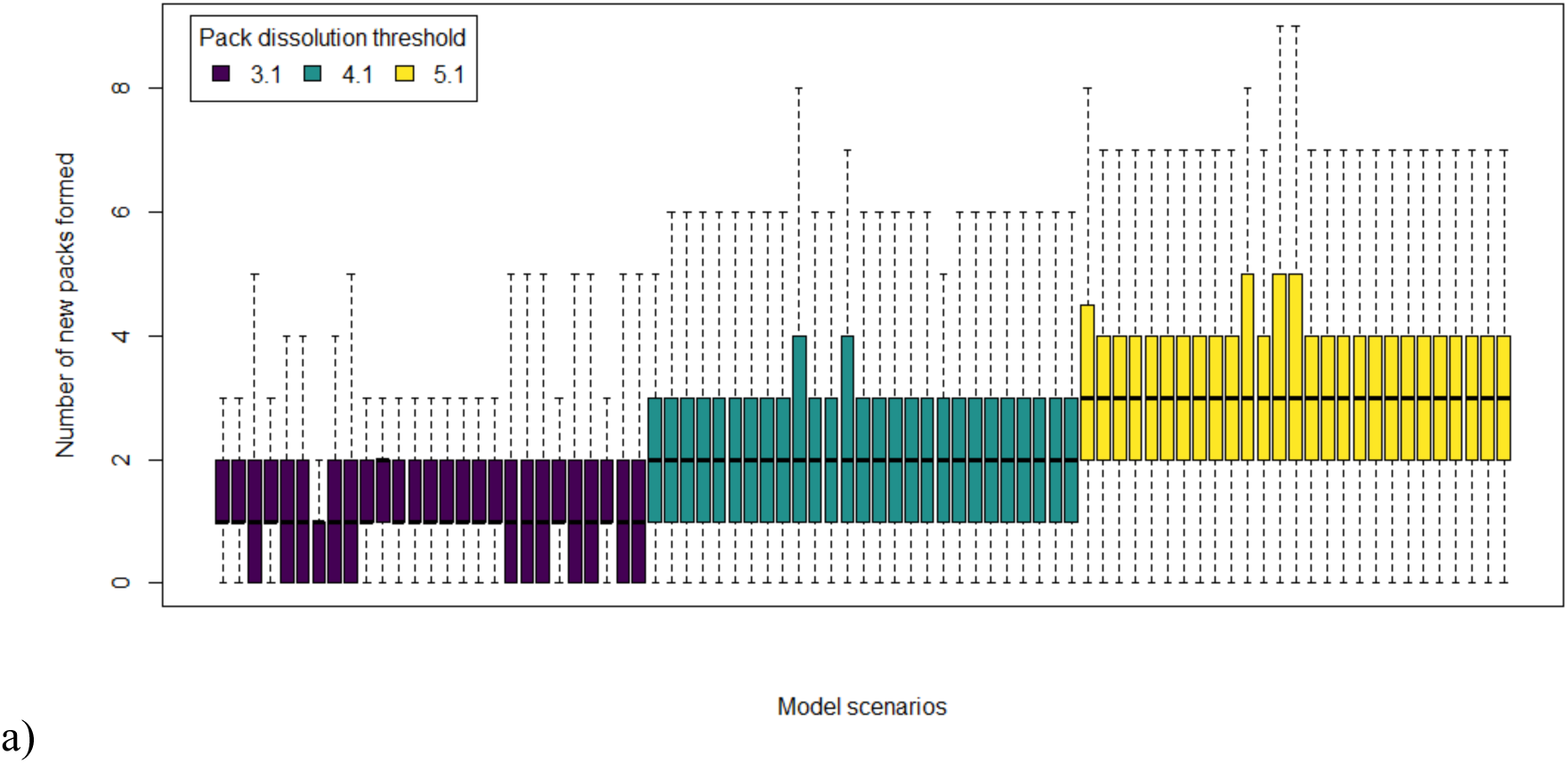
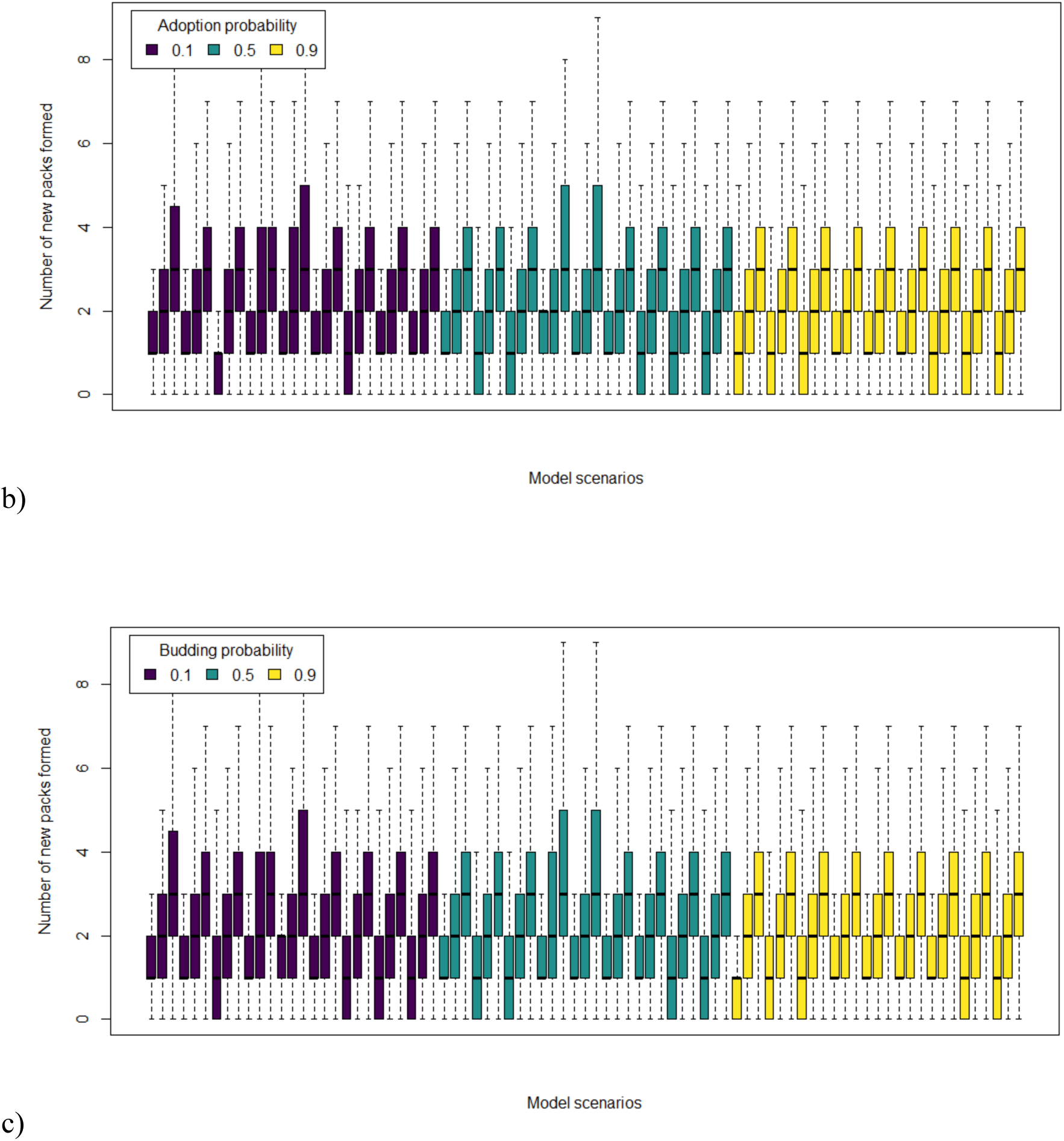
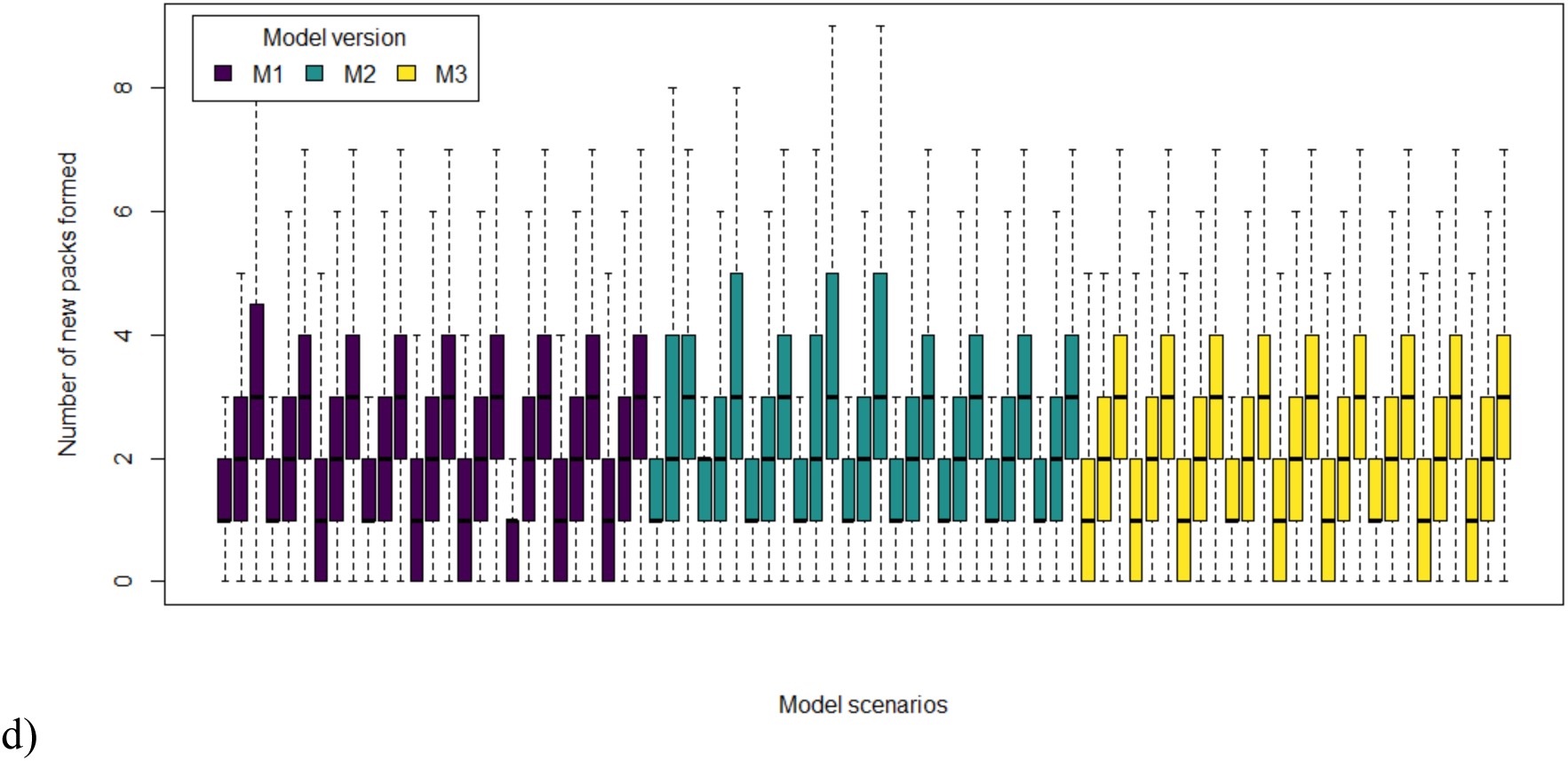
Number of new packs created during the last simulated year for each model scenario.

**Figure C4:**
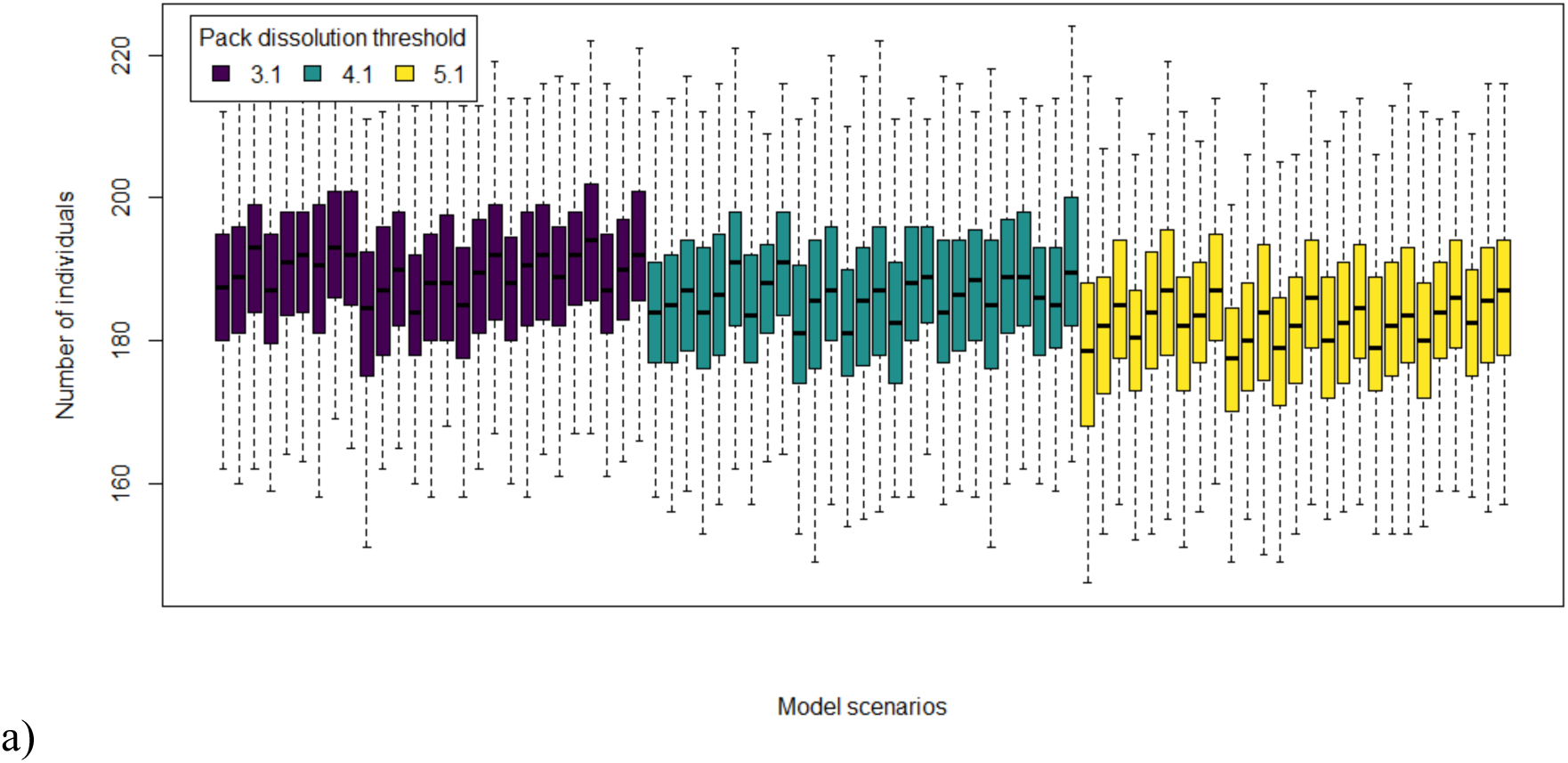
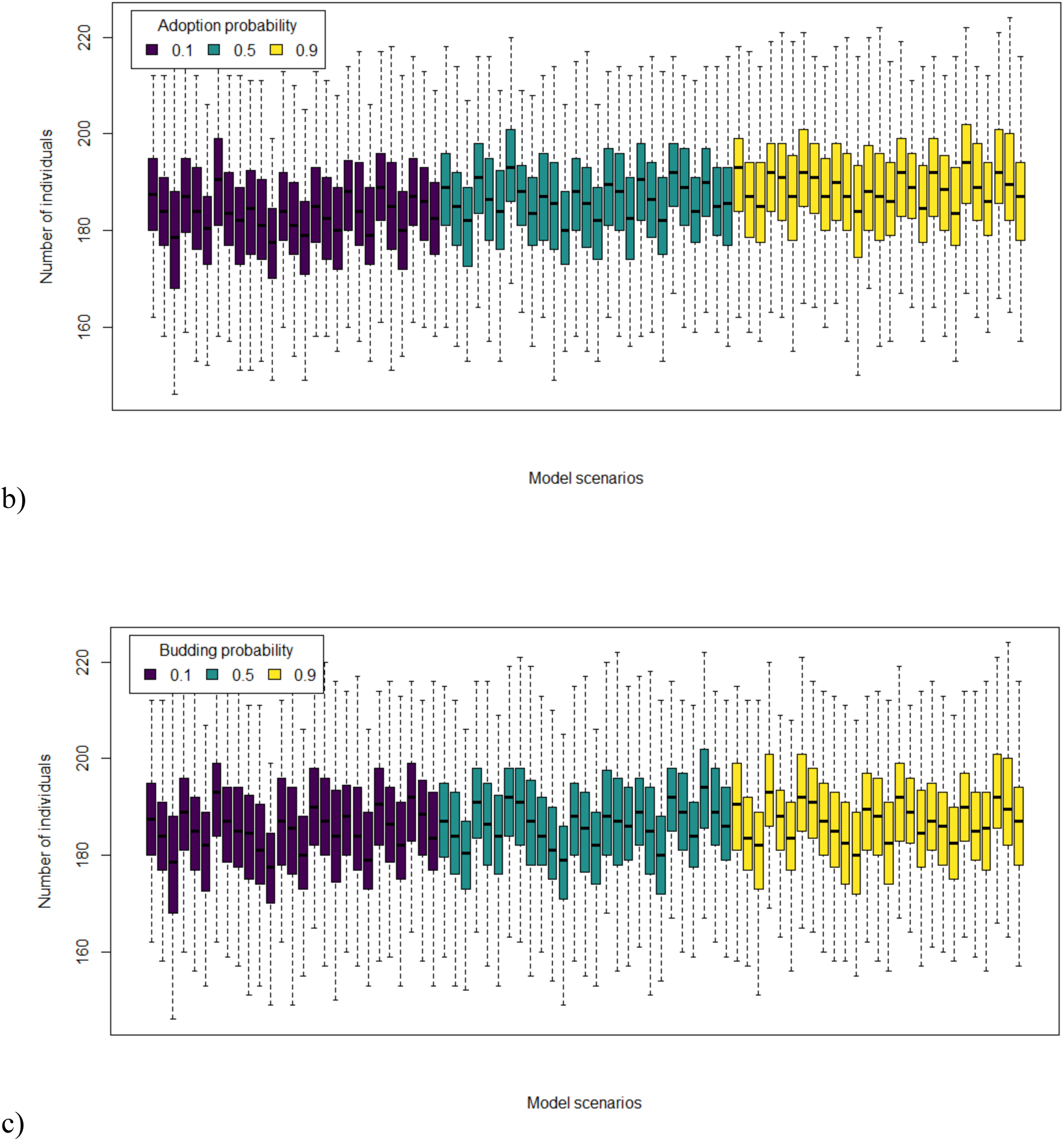
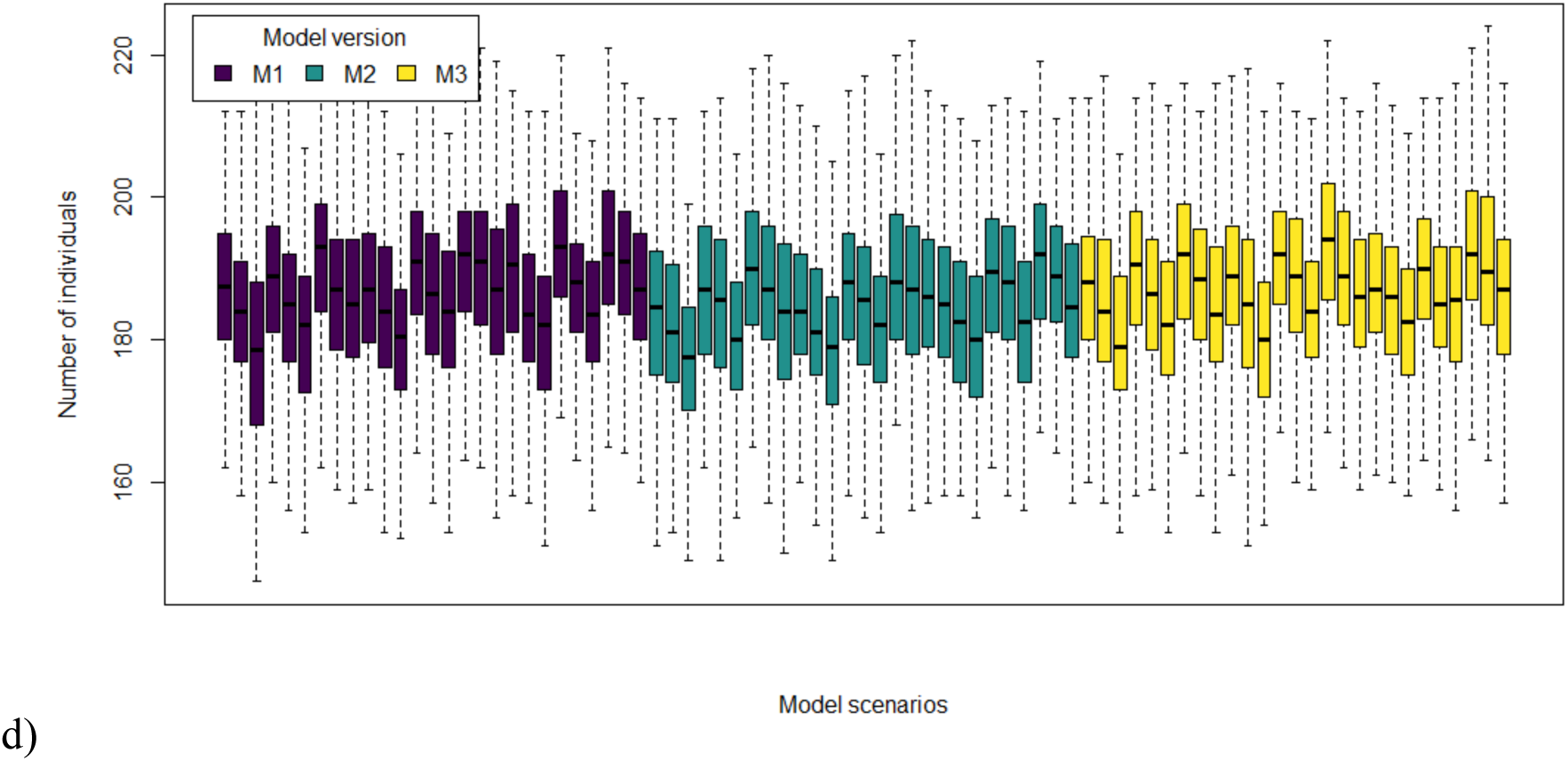
Total number of individuals in the population at the end of last simulated year for each model scenario.

**Figure C5:**
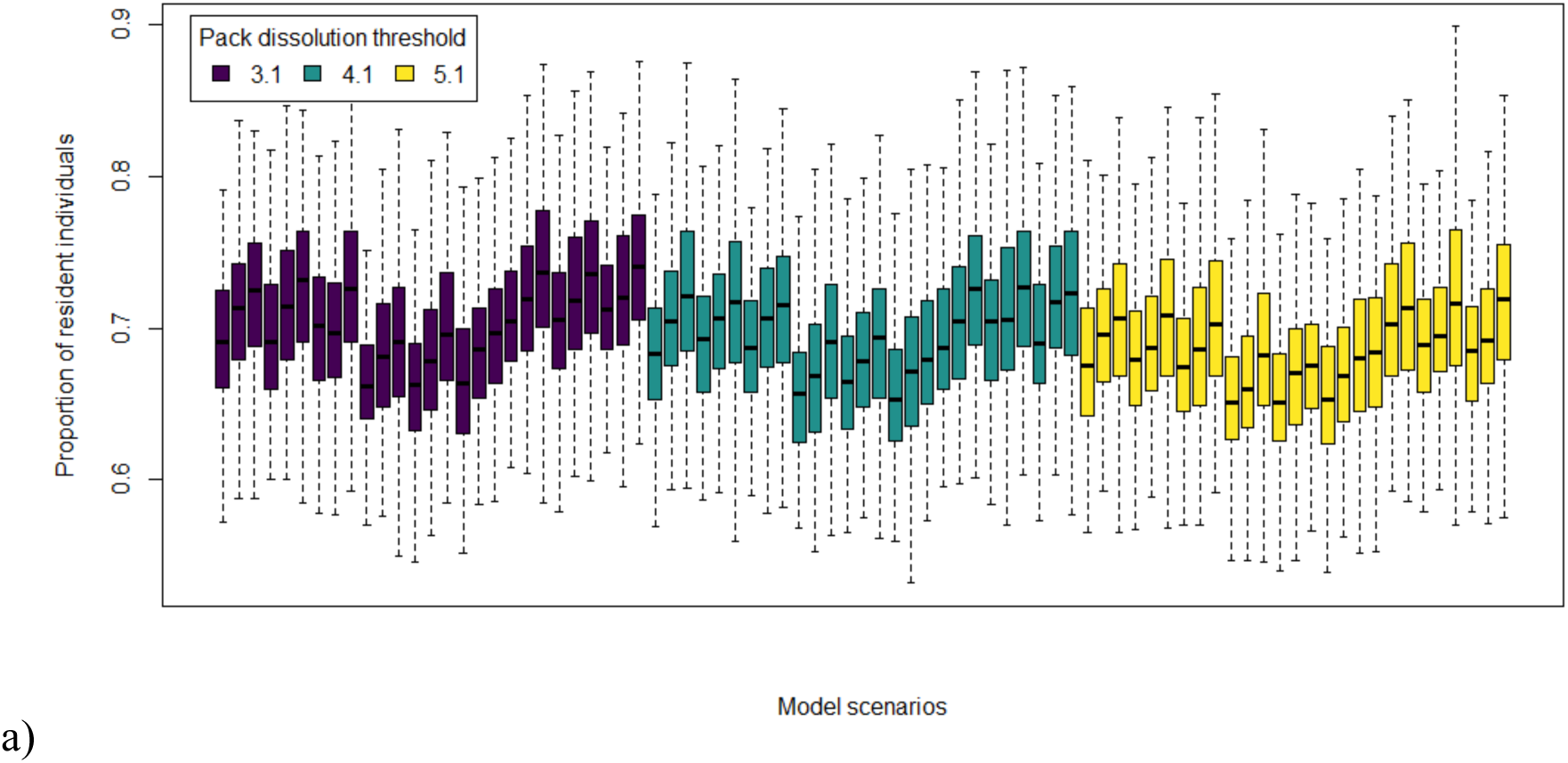
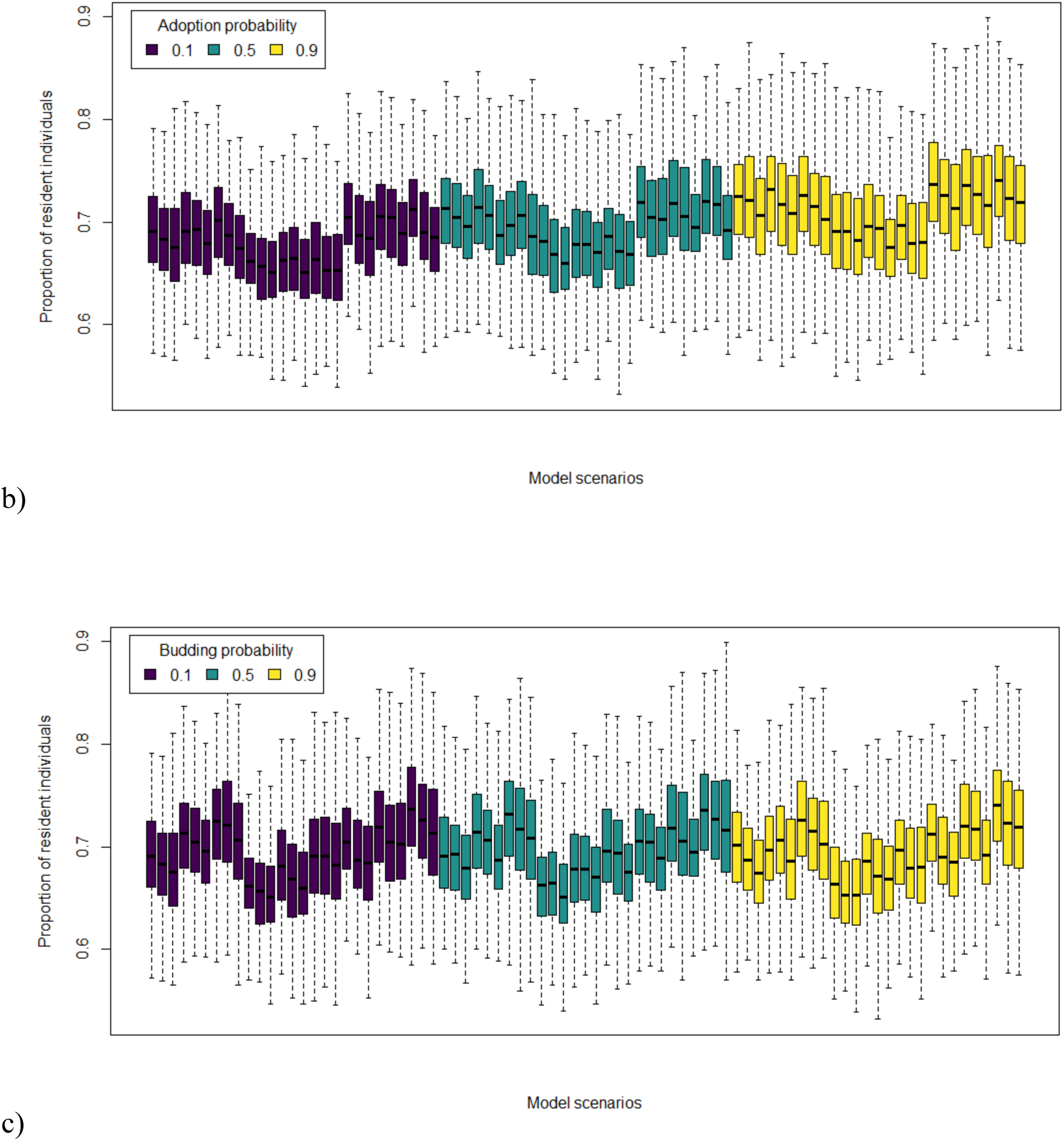
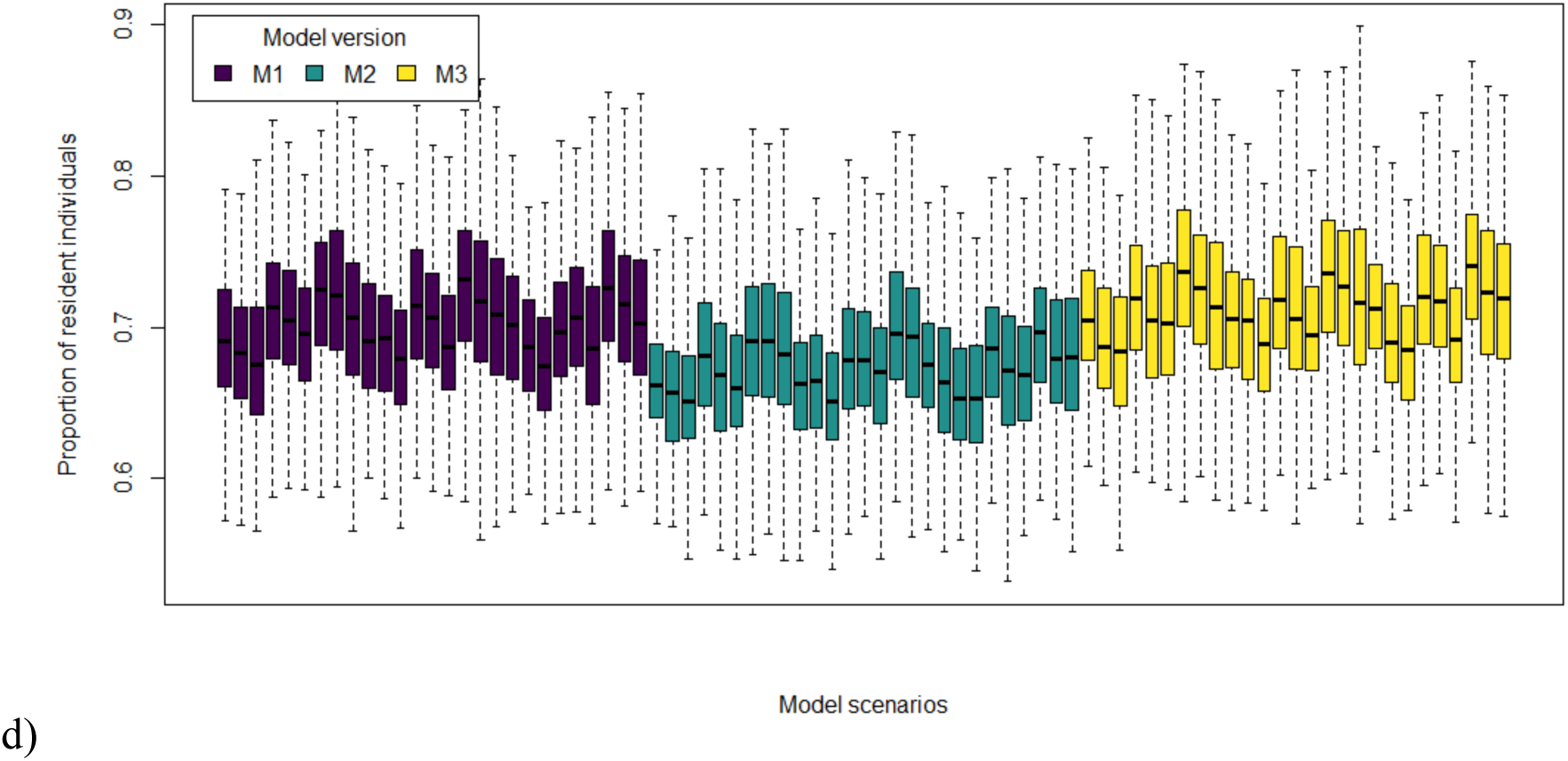
Proportion of resident individuals in the population at the end of last simulated year for each model scenario.

**Figure C6:**
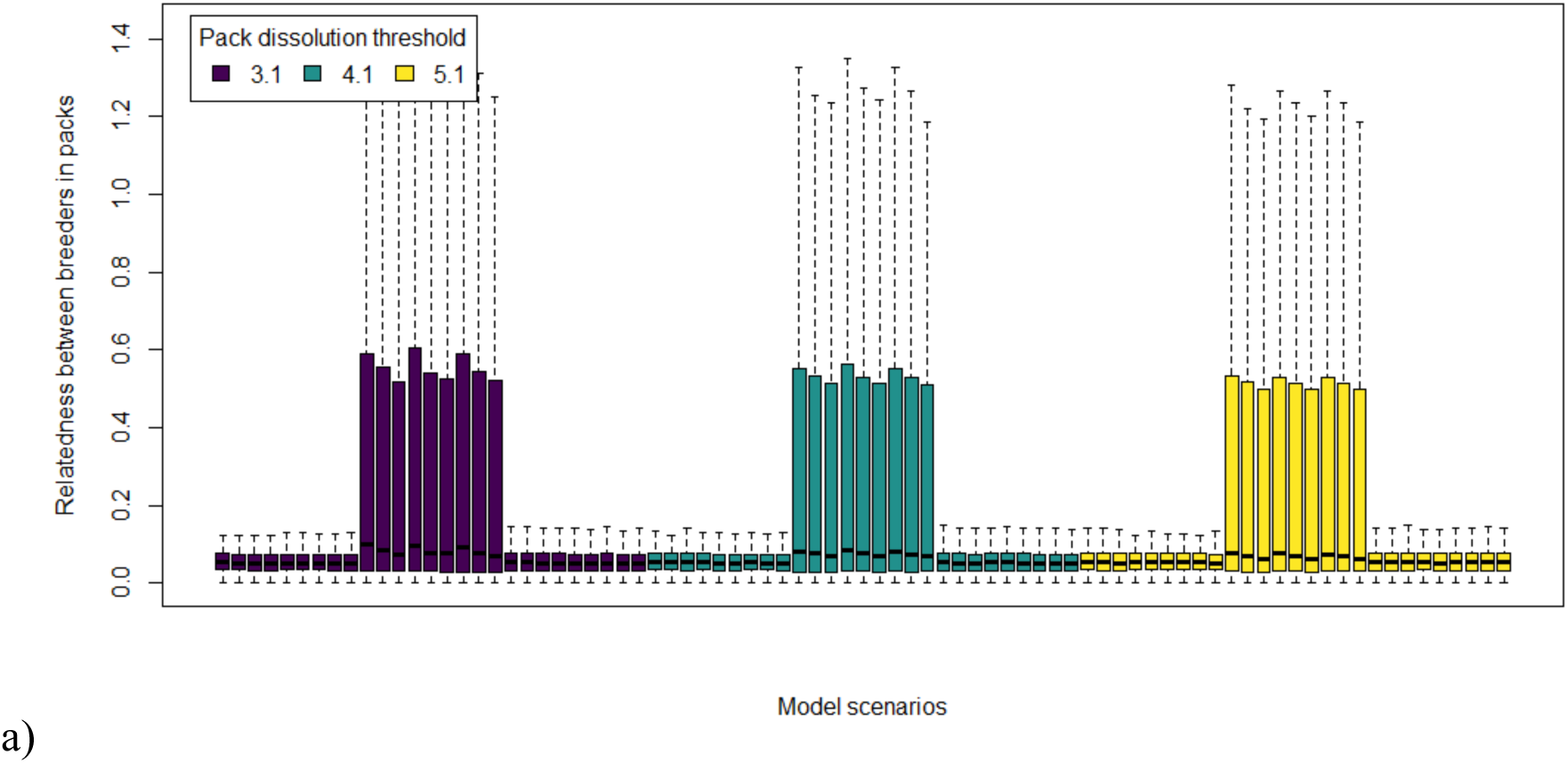
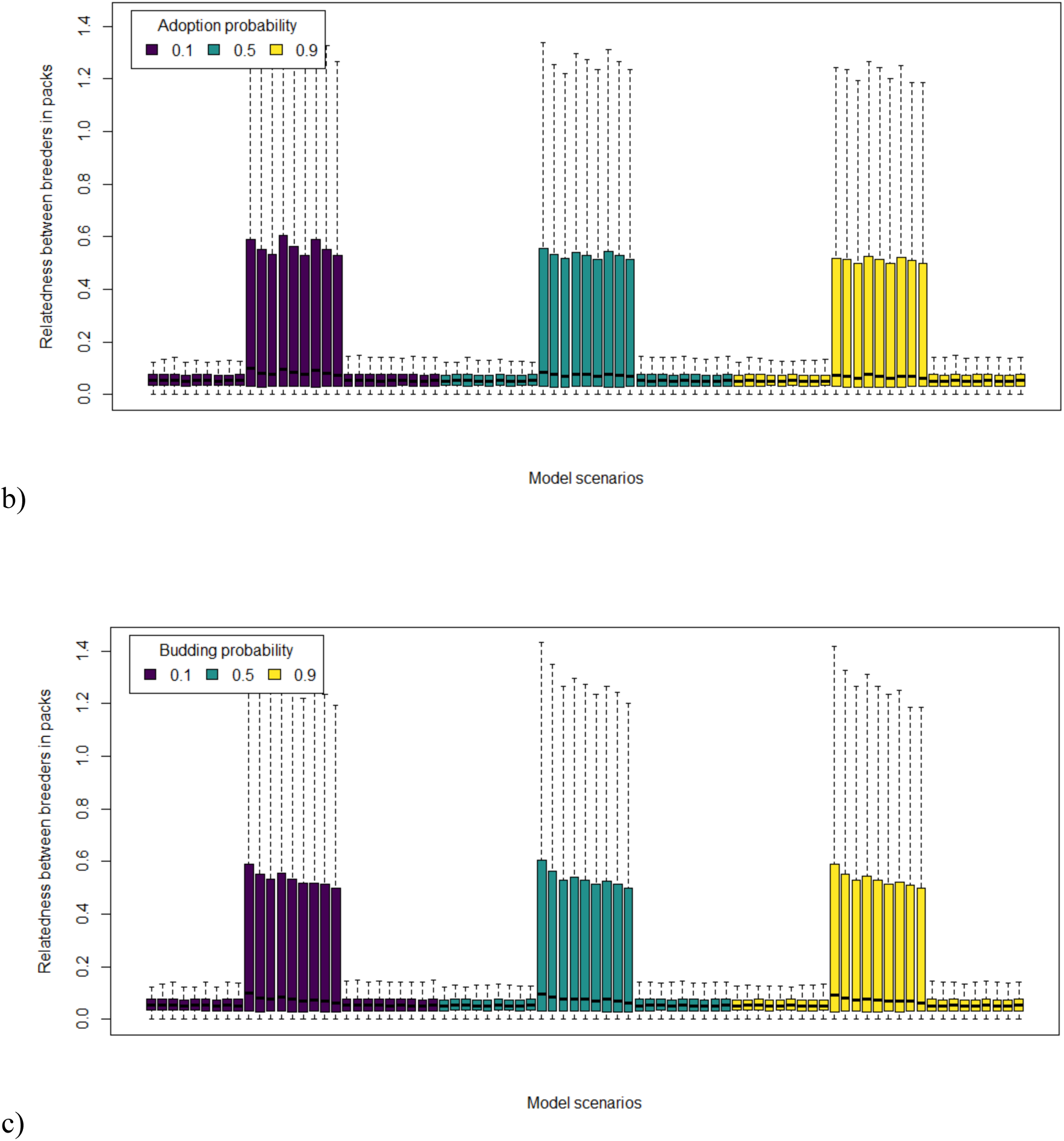
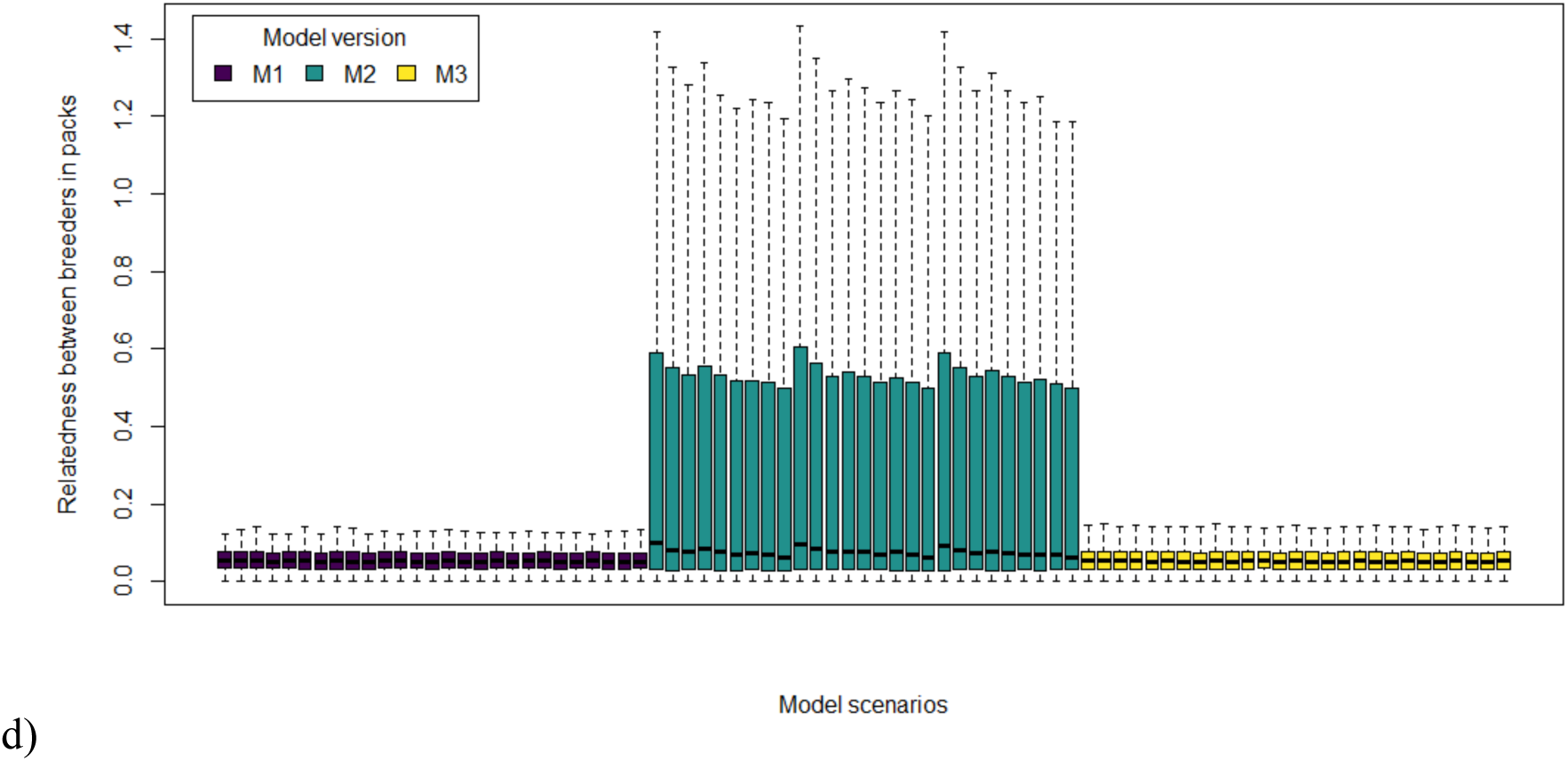
Relatedness value between the male and female in breeding pairs at the end of last simulated year for each model scenario.

## Appendix D

### File: sensitivityAnalysisResults.xlsx

Complete results of the sensitivity analysis. The first line of the table is the name of the simulation run: SA0 for the reference model and the runs SA1 to SA36 are the runs similar to SA0 where one parameter of the model was modified, one at a time, with its value either decreased or increased by 5 %. SA0 is the model version M1 (Fig. 1, main text), with the value 4.1, 0.5, and 0.5 for the parameters of sub-models pack dissolution, adoption and establishment by budding respectively (model scenario S14 from Appendix B), and the parameter values from the up-to-date literature for the other sub-models (Table 1, main text). The second line of the table informs which parameter was modified in the sensitivity analysis run and the following line gives the value used for this parameter. Then, the six following line are the six model outputs: the year at which populations reached equilibrium density, the number of packs with a breeding pair, the number of new packs created, the number of individuals, the proportion of resident individuals and the relatedness between the individuals in breeding pairs. The result values are the mean values over the 200 simulation replicates for each run. The column “SA0 [- 20 %; + 20 %]” presents the results for the run with reference model with the range – 20 % and + 20 % of the result values. Then, table cells are the mean values of the model outputs obtained with the runs SA1 to SA36. Dark orange cells are model results outside of the reference range of M0 results [- 20 %; + 20 %], light orange cells are the lowest and highest values for the model outputs.

